# A conserved mechanism for targeted cohesin loading organises specialised chromosomal domains

**DOI:** 10.64898/2026.07.21.739850

**Authors:** Hollie J. Rowlands, Meg Peyton-Jones, Daniel Robertson, Shaun Webb, Christos Spanos, Adele L. Marston

**Affiliations:** Centre for Cell Biology, Institute of Cell Biology, School of Biological Sciences, Michael Swann Building, Max Born Crescent, Edinburgh, EH9 3BF UK; Discovery Research Platform for Hidden Cell Biology, School of Biological Sciences, University of Edinburgh, Edinburgh EH9 3BF, UK

**Keywords:** Cohesin, *S. pombe*, chromosome segregation, genome organisation, rDNA, pericentromeres

## Abstract

Cohesin organises chromosomes through loop extrusion, yet how cohesin loading is targeted to specific genomic regions to shape chromosome architecture remains poorly understood. Here, we identify a conserved mechanism that directs cohesin loading to specialised chromosomal domains in *Schizosaccharomyces pombe*. Mutation of a conserved interaction surface in the cohesin loader Scc4/Ssl3 abolishes cohesin enrichment at centromeres and ribosomal DNA while largely preserving cohesin levels along chromosome arms, demonstrating that domain-specific accumulation depends on targeted recruitment rather than global loading capacity. We further identify the nucleolar protein Dnt1 as a receptor for cohesin loading at rDNA and show that it is required for proper organisation of this domain. High-resolution MicroC-XL maps reveal that targeted cohesin loading positions loop-based interactions within pericentromeres and rDNA without disrupting larger-scale domain boundaries. Together, these findings show that domain-specific receptors acting through a conserved loader interface direct cohesin positioning and identify targeted loading as a conserved organising principle for specialised chromosomal domains.

## Introduction

The <u>S</u>tructural <u>M</u>aintenance of <u>C</u>hromosomes (SMC) protein family dynamically organise genomes to control chromosome segregation, DNA repair and gene expression (Yatskevich *et al*, 2019; Hoencamp & Rowland, 2023). Their fundamental property, conserved from prokaryotes to eukaryotes, is the ability to form DNA loops, These loops are thought to arise through loop extrusion, a process that progressively enlarges DNA loops and organises chromosomes into domains of enriched interactions (Fudenberg *et al*, 2016; Nasmyth, 2001). Accordingly, chromosome capture methods *in vivo* and single molecule studies *in vitro* support cohesin-dependent loop extrusion as a fundamental mechanism of chromosome organisation (Rutkauskas & Kim, 2025; Davidson & Peters, 2021). Because loop extrusion initiates at sites of cohesin loading, mechanisms that target the cohesin loader to specific chromosomal locations are likely to play an important role in determining how chromosome architecture is organised. While loop extrusion resulting in the juxtaposition of two segments along one molecule of DNA appears to be a feature of all SMC complexes, eukaryotic cohesin has uniquely acquired an additional property: the ability to hold two sister DNA molecules together. This sister chromatid cohesion is generated through a poorly understood mechanism that is coupled to replication fork passage, resulting in entrapment of the newly duplicated DNA strands within the ring-shaped cohesin complex (Murayama, 2025). Cohesin is comprised of two SMC proteins, Smc1 and Smc3, which form antiparallel coiled-coils that heterodimerise at their hinge regions, and whose ATPase head domains are connected by the alpha-kleisin Scc1/Rad21. Both cohesin loading onto chromosomes and loop extrusion rely on a separate conserved complex, known as Scc2-Scc4 in *Saccharomyces cerevisiae*, Mis4-Ssl3 in *Schizosaccharomyces pombe* or NIPBL-MAU2 in humans (Yatskevich *et al*, 2019; Hoencamp & Rowland, 2023).

Although SMC complexes act broadly across the genome they are also selectively enriched at specialised chromosomal sites such as pericentromeres and the rDNA. How SMC complexes are specifically targeted to these regions and how such targeted loading contributes to chromosomal architecture remain central questions. To date, *S. cerevisiae* pericentromeres provide the only system in which targeted cohesin loading has been mechanistically defined. The ∼125 bp centromere assembles the kinetochore (McAinsh & Marston, 2022), which not only anchors microtubules but also recruits cohesin (Megee *et al*, 1999; Weber *et al*, 2004; Eckert *et al*, 2007; Ng *et al*, 2009; Fernius *et al*, 2013; Fernius & Marston, 2009). This process depends on the Dbf4-dependent kinase (DDK), which phosphorylates the N-terminal region of the inner kinetochore subunit Ctf19^CENP-P^, creating a docking site for a conserved patch on the Scc4 component of the cohesin loader (Hinshaw *et al*, 2017, 2015). Kinetochore-directed cohesin loading shapes pericentromeres into loops and is essential for establishing robust cohesion at loop bases to resist spindle forces and ensure accurate chromosome segregation (Paldi *et al*, 2020).

Despite this insight, *S. cerevisiae* pericentromeres are atypical. In most eukaryotes, centromeres are epigenetically specified and embedded within heterochromatic domains, but they are nonetheless enriched in cohesin (Sacristan *et al*, 2024; Bernard *et al*, 2001; Schmidt *et al*, 2009). In *S. pombe*, pericentromeric cohesin accumulation depends on the heterochromatin protein Swi6 (HP1 ortholog), as well as the DDK kinase Hsk1–Dfp1 (Bailis *et al*, 2003; Nonaka *et al*, 2002; Bernard *et al*, 2001), suggesting heterochromatin-directed mechanisms. However, whether comparable targeted loading mechanisms operate beyond *S. cerevisiae* remains unknown.

The ribosomal DNA (rDNA) locus also requires specialised spatial organisation within the nucleolus to maintain genome stability and support ribosome biogenesis. Comprising hundreds of tandemly repeated transcription units, rDNA arrays pose unique topological challenges due to high RNA polymerase I activity and susceptibility to recombination and replication stress (Hori *et al*, 2023). Both cohesin and condensin safeguard rDNA integrity by regulating DNA double strand break repair (Huang *et al*, 2006; Kobayashi *et al*, 2004; Kobayashi & Ganley, 2005; Marnef *et al*, 2019; Li *et al*, 2014) and promote nucleolar condensation and segregation (Lavoie *et al*, 2004, 2000, 2002; Freeman *et al*, 2000; D’Ambrosio *et al*, 2008). Cohesin further supports rDNA replication and ribosome biogenesis in yeast and humans (Lu *et al*, 2014; Harris *et al*, 2014; Bose *et al*, 2012). In *S. cerevisiae,* condensin recruitment to the rDNA requires the Replication Fork Barrier (RFB) protein, Fob1, which binds directly to the InterGenic Spacer region (IGS1) of the rDNA repeat and assembles a nucleolar protein complex, comprising cohibin (Lrs4-Csm1) and Tof2 (Johzuka *et al*, 2006; Huang *et al*, 2006). Condensin recruitment to the *S. cerevisiae* mating type locus has similar requirements (Dinda *et al*, 2023). We recently found that condensin loading is targeted to specific sites via direct interaction of a conserved pocket on the condensin HAWK subunit, Ycg1, with a short linear “CR1” motif in chromosomal receptors, among them Lrs4 in the rDNA (Wang *et al*, 2025). Cohesin associates with a distinct region between the rDNA repeats, termed the non-transcribed spacer (NTS2) (Laloraya *et al*, 2000), but the mechanisms involved in its recruitment are unclear.

The emerging picture is that SMC complexes use conserved surfaces on their regulatory subunits to interact with short-linear motifs on chromosome-located receptors, thereby directing their activities to specific genomic sites with high structural needs. In parallel, a non-targeted loading mechanism ensures SMC distribution across the genome. We hypothesised that disrupting these conserved interaction surfaces would selectively impair targeted cohesin loading and local chromatin architecture while leaving global genome organisation largely unaffected. To test this, we mutated the Ssl3 patch of the cohesin loader in fission yeast *S. pombe*, an organism whose heterochromatic pericentromeres resemble those of higher eukaryotes yet remain experimentally tractable. We identify pericentromeres and rDNA as sites dependent on targeted cohesin loading via the conserved Ssl3 patch and reveal the nucleolar protein Dnt1 as a receptor for cohesin loading in the rDNA. Using MicroC-XL, we show that targeted cohesin loading positions interactions within pericentromeres and rDNA, while global chromosome structure is largely independent of targeted cohesin loading. Overall, our findings reveal a conserved mechanism for targeted cohesin loading and demonstrate its importance for the organisation of functional chromosomal domains.

## Results

### Domain-specific localisation of cohesin and its loader Mis4 in *S. pombe*

Previous work suggested that cohesin loading and retention are differentially regulated across the genome (Schmidt *et al*, 2009). As a baseline for understanding how cohesin is targeted in *S. pombe*, we used calibrated ChIP-Seq in cycling *S. pombe* cells, the majority of which are in G2, to compare the genome-wide localisation of cohesin Rad21^Scc1^ and its loader/activator Mis4^Scc2^. To account for differences in rDNA copy number between strains, we routinely mapped our ChIP-seq data to a custom genome file containing only the rDNA sequence to produce ChIP-seq profiles that are pileups of rDNA data normalised to the number of rDNA sequences in our Input controls. This revealed enrichment of both Rad21^Scc1^ and Mis4^Scc2^ at pericentromeres and the rDNA locus, with smaller peaks along chromosome arms (Fig. 1A). Despite overall co-enrichment, Rad21^Scc1^ and Mis4^Scc2^ displayed distinct distributions within pericentromere regions (Fig. 1A,B). Rad21^Scc1^ was most enriched over the heterochromatic outer repeats (*otr*) and the *otr*-adjacent segment of the inner repeats (*imr*) that extends to the first tRNA gene; however, Rad21^Scc1^ is relatively depleted from the rest of the inner repeats and central core (*cc*), the primary site of kinetochore assembly. In contrast, Mis4^Scc2^ was most enriched across the core centromere and inner repeats, with lower levels on the outer repeats. Both cohesin and Mis4 are highly enriched at centromeric tRNAs (Fig. 1B). Given that Mis4^Scc2^ associates with actively loading or extruding cohesin (Fernius *et al*, 2013; Petela *et al*, 2018), this suggests a dynamic cohesin pool at the core and inner repeat regions. Enrichment of Rad21 at the heterochromatic outer repeats is consistent with a stabilised cohesin pool, in agreement with the requirement for the heterochromatin protein Swi6 for robust pericentromeric cohesion and accurate chromosome segregation (Bernard *et al*, 2001; Nonaka *et al*, 2002). At the rDNA locus, Mis4^Scc2^ and Rad21^Scc1^ showed highly similar profiles, with strongest signal at the non-transcribed spacer (NTS) region and smaller peaks within the rRNA transcription unit (Fig. 1C). Along chromosome arms, Rad21 peaks frequently occurred between convergent genes with minimal Mis4^Scc2^ enrichment, whereas tRNA genes showed enrichment for both proteins (Fig. 1D), which could indicate sites of cohesive and looping cohesin, respectively.

**Figure 1.**
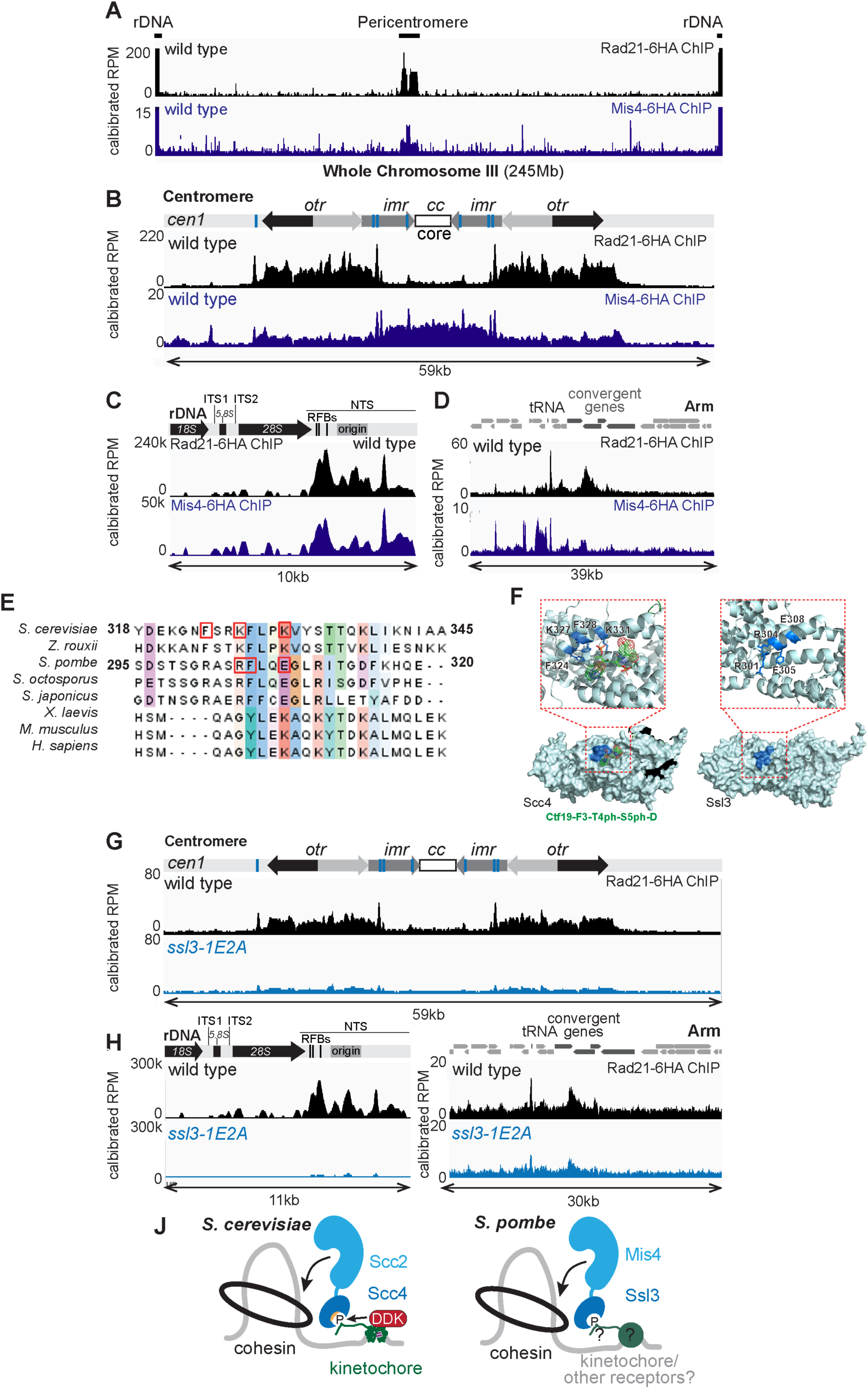
A conserved patch on Ssl3 targets cohesin loading to pericentromeres and rDNA. (A-D) Distinct patterns of cohesin (Rad21) and its loader/regulator (Mis4) on chromosomes in wild type cells. ChIP-Seq of Rad21-6HA (upper tracks, black; AMfy638) and Mis4-6HA (lower tracks, grey; AMfy659). View of whole chromosome III (A) and zoomed in pericentromeres (B), rDNA (C) and a region on the arm of chromosome I, including a tRNA site and two pairs of convergent genes (D). (E) Alignment of Scc4/Ssl3 homologs. (F) Comparison of Scc4 crystal structure (Hinshaw *et al*, 2017) with alphafold models of Scc4/Ssl3 patch. (G-I) Rad21 ChIP-Seq in wild type and *ssl3-1E2A* for centromere (G), rDNA (H) and arm (I) regions. *S. pombe* strains used are AMfy2157 (wild type) and AMfy2202 (*ssl3-1E2A*). (J) Models for targeted loading of cohesin in *S. cerevisiae* and *S. pombe*. See also Fig S1.

To test whether these domain-specific patterns reflect distinct recruitment mechanisms, we examined genetic dependencies. Deletion of the H3K9 methyltransferase Clr4 abolished Rad21 enrichment across the pericentromeric outer repeats, had modest effects on the distribution at the rDNA locus and no effect along chromosome arms (Fig. S1A-C). Conversely, kinetochore inactivation using the temperature-sensitive *fta2-291* allele (Kerres *et al*, 2006) eliminated Mis4 enrichment at the central core and inner repeats. However, Rad21 was modestly increased across the outer repeats and at tRNAs within the pericentromeric region in *fta2-291* cells, including those flanking the central core, potentially due to redistribution (Fig. S1A). This indicates that Mis4 and Rad21 enrichment at tRNAs and heterochromatic repeats are independent of the kinetochore. Together, these analyses identify pericentromeres and the rDNA regions as specialised chromosomal domains with distinctive cohesin-based organisation, indicating that local chromosomal features impose domain-specific modes of cohesin regulation.

To address the role of DDK in cohesin localisation we also examined Rad21 enrichment in the presence of a C-terminal truncation of Dfp1 (*dfp1-(1-376)*), which is defective in Swi6 interaction and was shown to cause cohesin loss at an integrated pericentromeric reporter by ChIP-PCR (Bailis *et al*, 2003). Interestingly, our results indicate that cohesin levels at pericentromeres, rDNA and on chromosomal arms in the *dfp1-(1-376)* mutant are comparable to wild type (Fig. S1D-F), likely reflecting the enhanced accuracy of calibrated ChIP-seq for examining chromosomal protein enrichment. We note, however, that the Dfp1 truncation still activates DDK, and thus this finding does not preclude a role for DDK in cohesin enrichment.

### A conserved patch on Ssl3 targets cohesin to centromeres and rDNA

To test whether cohesin enrichment at specialised chromosomal domains depends on targeted loading, we examined the role of a conserved interaction surface in the cohesin loader Ssl3^Scc4^. In *S. cerevisiae*, this surface docks onto kinetochores to direct cohesin loading and pericentromere organisation (Hinshaw *et al*, 2017, 2015). Guided by sequence alignment (Fig. 1E) and structural comparison of *S. cerevisiae* Scc4 (Hinshaw *et al*, 2017, 2015) with an Alphafold model of Ssl3 (Fig. 1F), we generated the *ssl3-R304E, F305A, E308A* mutant (*ssl3-1E2A*), which we predicted should disrupt targeted cohesin loading. Indeed, in *ssl3-1E2A* cells, specific cohesin enrichment at centromeres, pericentromeres and rDNA, was largely lost, while cohesin association along chromosomal arms was only slightly reduced (Fig. 1G-I). These results indicate that mutation of the conserved Ssl3 patch has a relatively limited effect on overall genomic cohesin levels but severely impairs enrichment at specialised chromosomal regions. We conclude that targeted cohesin enrichment on *S. pombe* chromosomes depends on the conserved interaction surface of the cohesin loader Ssl3 (Fig. 1J).

### Bypassing Ssl3-dependent Mis4 activation does not restore targeted cohesin loading

To confirm that cohesin loss at centromeres and rDNA in the *ssl3-1E2A* mutant reflects a defect in targeted recruitment rather than general loading activity, we sought to overcome the essential requirement for Ssl3 in cohesin loading (Bernard *et al*, 2006). In *S. cerevisiae*, the gain-of-function *scc2E822K L937F* allele enhances cohesin loading and bypasses the requirement for Scc4 for cohesin loading on chromosome arms, yet pericentromeric enrichment remains Scc4-dependent (Petela *et al*, 2018). The corresponding residues are highly conserved (Fig. S2A) and introduction of equivalent mutations (E907K L1027F, henceforth *mis4-1K1F*) into *S. pombe mis4* rescued the temperature-sensitive growth defect of the *ssl3-29* allele at 32°C (Fig. 2A). Importantly, the *mis4-1K1F* allele did not increase chromosomal cohesin levels in otherwise wild type cells but restored cohesin loading genome-wide in the *ssl3-29* mutant at the restrictive temperature (Fig. S2B-D). This indicates that *mis4-1K1F* bypasses the requirement for Ssl3 in promoting Mis4-dependent loading. However, because *ssl3-29* retains partial cohesin loading activity even at the restrictive temperature, this system could not resolve whether Ssl3 is specifically required for targeted cohesin recruitment. We therefore combined *mis4-1K1F* with the targeting-defective *ssl3-1E2A* mutant (Fig. 2B). Although *mis4-1K1F* restored cohesin association with chromosome arms in *ssl3-1E2A* background to near wild-type levels, cohesin enrichment at centromeres and pericentromeres was not recovered, and only a marginal increase in cohesin was observed at rDNA (Fig. 2C-E). Consistently, pileups show that *mis4-1K1F* restores cohesin peaks at arm features (long terminal repeats, LTR retrotransposons and tRNAs) but not centromeric tRNAs (Fig. S2E). Thus, removing the dependence on Ssl3 for Mis4-dependent cohesin loading is insufficient to restore cohesin accumulation at specialised chromosomal domains in the absence of the conserved Ssl3 targeting surface.

**Figure 2.**
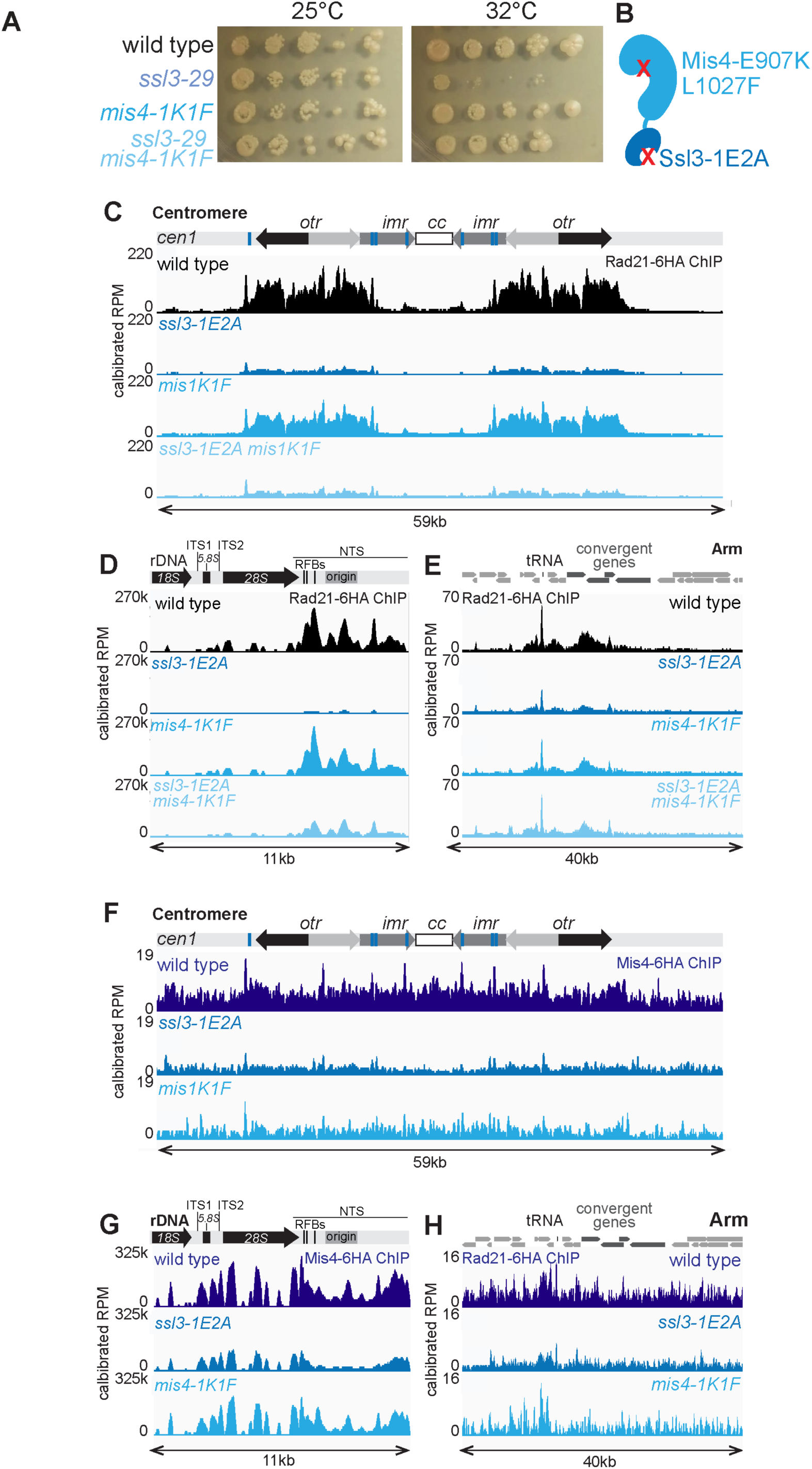
Ssl3-dependent Mis4 activation and targeting are genetically separable. (A) Mis4-E907K L1027F (Mis4-1E1F) rescues growth of *ssl3-29* at the restrictive temperature. Spotting assay comparing growth of wild type (AMfy2157), *ssl3-29* (AMfy2158), *mis4-E907K L1027F* (AMfy2163) and *ssl3-29 mis4-E907K L1027F* (AMfy2162) strains at the indicated temperatures. (B) Schematic of Mis4-Ssl3 showing the locations of the indicated mutations. (C-E) Mis4-1E1F fails to rescue cohesin association with pericentromeres and rDNA in *ssl3-1E2A* cells. Rad21-6HA ChIP-Seq profiles of centromeres (C), rDNA (D) and a representative chromosomal arm region (E) for wild type (AMfy2157), *mis4-E907K L1027F* (AMfy2164), *ssl3-1E2A* (AMfy2202), *ssl3-1E2A mis4-E907K L1027F* (AMfy2196). (F-H) Ssl3 patch mutations and Mis4 bypass mutations cause Mis4 mis-localisation at centromeres and the rDNA. Mis4-6HA ChIP-Seq profiles of centromeres (F), rDNA (G) and a representative chromosomal arm region (H) for wild type (AMfy2282), *ssl3-1E2A* (AMfy2283), *mis4-E907K L1027F* (AMfy2286). See also Fig S2.

To determine how *mis4-1K1F* and *ssl3-1E2A* affect cohesin loader distribution, we performed Mis4 ChIP-seq. Because our original Mis4-6HA fusion was inviable in these mutant backgrounds, we used an alternative 6HA linker that retained viability, enabling ChIP-seq, albeit with low efficiency. In *ssl3-1E2A*, Mis4 association was almost completely lost from centromeres, with the strongest reductions across the core centromere and inner repeats (Fig. 2F). Mis4 occupancy was also reduced across the rDNA (Fig. 2G), including loss of the prominent NTS peaks, consistent with the severe depletion of cohesin at this locus (Fig. 1G), while Mis4 distribution was modestly reduced on chromosome arms in *ssl3-1E2A* cells (Fig. 2H). These findings suggest that the Ssl3 patch targets the Mis4/Ssl3 loader to specialised chromosomal domains. Examination of Mis4-1K1F localisation at centromeres revealed reduced occupancy along with a more even distribution and fewer distinct peaks (Fig. 2F). In contrast, Mis4-1K1F binding at the rDNA was only modestly reduced; however, its distribution was altered, resulting in relatively higher association at the RFBs (Fig 2G). On chromosome arms, Mis4 occupancy was largely retained, although subtle shifts in peak position were apparent (Fig. 2H). This is consistent with Mis4-1K1F becoming less constrained by Ssl3-dependent positioning, allowing cohesin loading to occur over a broader set of sites. However, because cohesin subsequently translocates from its loading sites to preferred chromosomal positions, these changes in loader distribution have little effect on the final cohesin occupancy profile (Fig. 2E). Taken together, these findings show that *mis4-1K1F* alters Mis4 distribution without substantially affecting cohesin positioning, even at specialised chromosomal domains.

### Dnt1 is a nucleolar receptor for cohesin recruitment to rDNA

To identify chromosomal receptors that recruit the cohesin loader, we performed Mis4-SZZ immunoprecipitation followed by mass spectrometry and compared to a no-tag control. Mis4 pulldown robustly recovered cohesin subunits (Psm1, Psm3 and Rad21) and known regulators of cohesin dynamics (Wpl1, Pef1 and Psy2 (Birot *et al*, 2020)) (Fig. 3A). In addition, nucleolar proteins were significantly enriched, with Dnt1 emerging as one of the most highly enriched and statistically significant interactors (Fig. 3A). Given the strong enrichment of cohesin at rDNA and the loss of rDNA targeting in the *ssl3-1E2A* patch mutant, we reasoned that Dnt1 might associate with the conserved patch on Ssl3 to mediate nucleolar cohesin recruitment. We therefore compared the interactome of wild-type Ssl3-SZZ with the Ssl3-1E2A-SZZ patch mutant. This revealed that wild type Ssl3 associated with nucleolar proteins including Dnt1, however these associations were lost in Ssl3-1E2A-SZZ pulldowns, despite retention of Mis4 binding (Fig. 3B, Fig S3A,B). Together, these observations indicate that mutation of the conserved Ssl3 patch does not destabilise the Ssl3-Mis4 complex, but rather disrupts its interaction with at least one potential receptor, the nucleolar protein, Dnt1.

**Figure 3.**
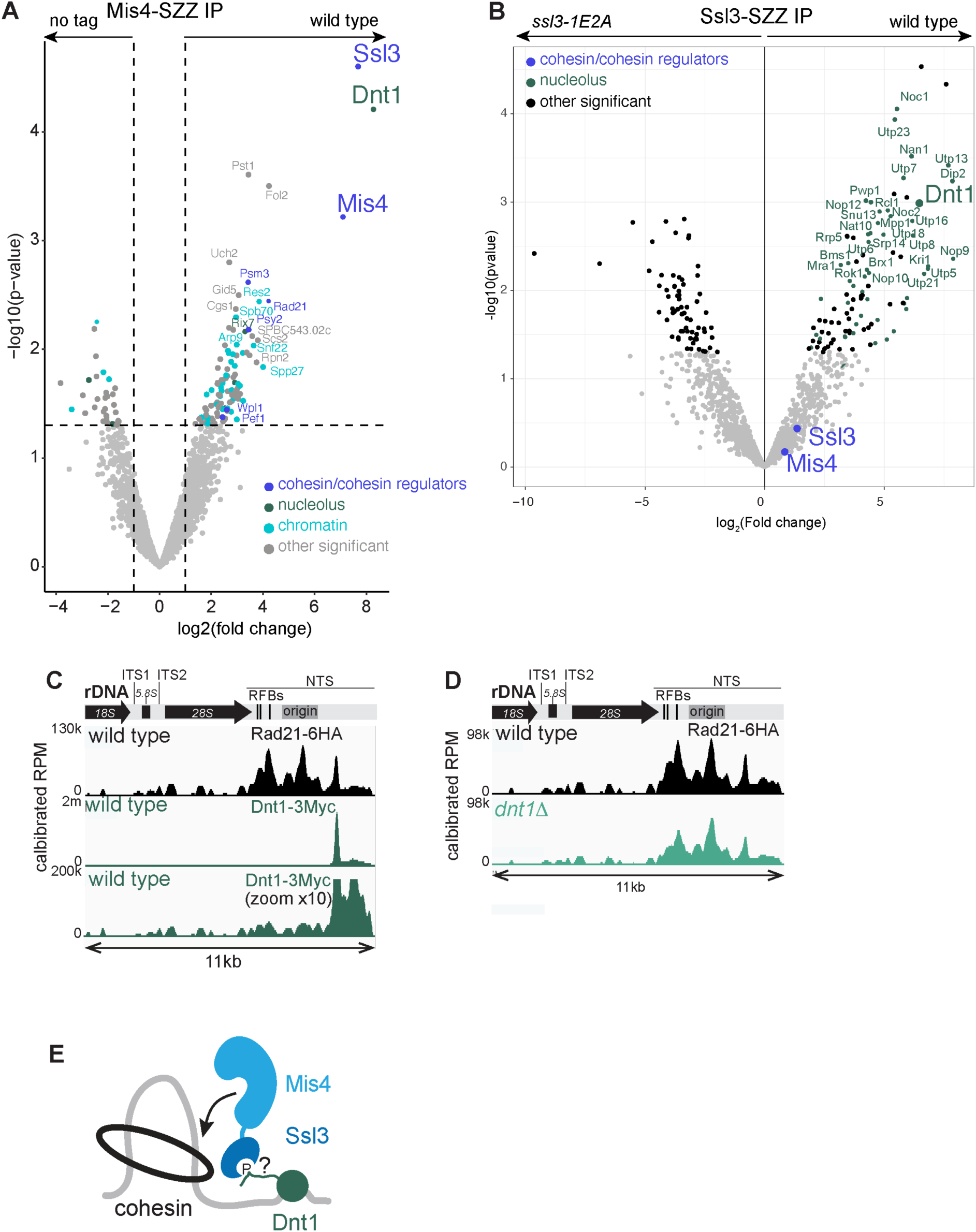
Dnt1 is a nucleolar receptor for targeted cohesin loading. (A) Dnt1 interacts with Mis4. Volcano plot comparison of proteins identified by mass spectrometry after immunoprecipitation of Mis4-SSZ compared to a no-tag control. Strains were AMfy207 (*mis4-SZZ*) and AMfy29 (*mis4+* untagged). (B) Interaction of Mis4/Ssl3 with nucleolar proteins and Dnt1 depends requires a functional Ssl3 patch. Volcano plot of proteins identified by mass spectrometry after immunoprecipitation of Ssl3-SZZ compared to Ssl3-1E1A-SZZ. Strains are AMfy205 (*ssl3-SZZ*) and AMfy2094 (*ssl3-1E2A-SZZ*). (C) Dnt1 co-localises with cohesin at the rDNA. ChIP-Seq of Rad21-6HA (upper tracks, black; AMfy638) and Dnt1-9Myc plus 10X zoom (lower tracks, green; AMfy2203). (D) Dnt1 recruits cohesin to the rDNA. Rad21-6HA ChIP seq in wild type (AMfy638) and *dnt1Δ* (AMfy1359). (E) Model for cohesin loading at rDNA via interaction of Ssl3 patch with Dnt1. See also Fig S3.

We hypothesised that docking of the conserved Ssl3 patch onto Dnt1 directs cohesin enrichment at the rDNA. ChIP–Seq of Dnt1-9Myc revealed enrichment primarily within the NTS region of the rDNA locus, overlapping with cohesin (Rad21) (Fig. 3C). Dnt1 exhibited a single dominant peak within the NTS that precisely coincided with one of the major cohesin peaks, while lower levels of Dnt1 were detectable at additional cohesin-enriched sites (Fig. 3C). Thus, although Dnt1 and cohesin profiles overlap, Dnt1 is concentrated at a discrete site in the rDNA.

To test whether Dnt1 is required for cohesin recruitment at the rDNA, we analysed the localisation of cohesin Rad21 by ChIP-Seq in *dnt1Δ* cells. Rad21 levels were depleted across the rDNA locus in *dnt1Δ* cells, indicating that Dnt1-dependent targeting of Mis4/Ssl3 to the NTS is necessary for proper cohesin recruitment and accumulation throughout the rDNA region (Fig. 3D). Interestingly, Rad21 enrichment at pericentromeres was modestly increased in *dnt1Δ* cells, while chromosomal arm levels were comparable to wild type (Fig. S3D,E). This is reminiscent of Rad21 enrichment at pericentromeres in *fta2-291* cells (Fig. S1A), further supporting the idea that cohesin is limiting, such that loss of spatial cohesin cues in one domain can lead to enhanced loading at another targeted loading site. We conclude that Dnt1 is a nucleolar receptor for Mis4/Ssl3 which enriches cohesin throughout the rDNA (Fig. 3E).

### Cohesin establishes local chromatin domains in *S. pombe*

The *S. pombe* genome is shaped by cohesin-dependent globules which form locally interacting domains and have been previously visualised using a low-resolution (10kb bin) Hi-C approach (Mizuguchi *et al*, 2014). To more precisely understand how cohesin shapes *S. pombe* chromosomes we employed MicroC-XL to create high resolution contact matrices, enabling visualisation of chromatin contacts at 100bp resolution. Consistent with previous data, we observe inter-centromere and inter-telomere interactions indicative of their clustering (Fig. S4A). Examination of a ∼1.2 Mb region of chromosome I, together with a zoomed-in ∼40 kb segment, confirmed that chromosome arms are organised into small interaction domains. Contact maps showed triangular domains of enriched local interactions but lacked prominent punctate interaction peaks (Fig. 4A,B). This pattern is consistent with local chromatin folding over a range of genomic distances (∼1-40kb) rather than stable interactions between defined genomic loci. Insulation score minima aligned with Rad21 ChIP-seq peaks, indicating that cohesin demarcates the boundaries of these local interaction domains (Fig. 4A,B). The contact probability curve and its derivative for chromosome arms (excluding the rDNA and pericentromeres) showed a smooth, near-linear decay with increasing genomic distance (Fig. 4C), suggesting that cohesin promotes interactions across a continuum of distances within these cohesin-defined boundaries, rather than forming stable loops of a characteristic size. To identify common elements at domain boundaries we generated pileups of our MicroC-XL data centred on annotated genome features. This analysis revealed that tRNA genes, promoters and long terminal repeats (LTR), and to a lesser extent 5’UTRs are present at the borders of chromosomal domains (Fig. 4D). We conclude that cohesin together with tRNAs, promoters and LTRs organises the *S. pombe* genome into small, locally-interacting, domains.

**Figure 4.**
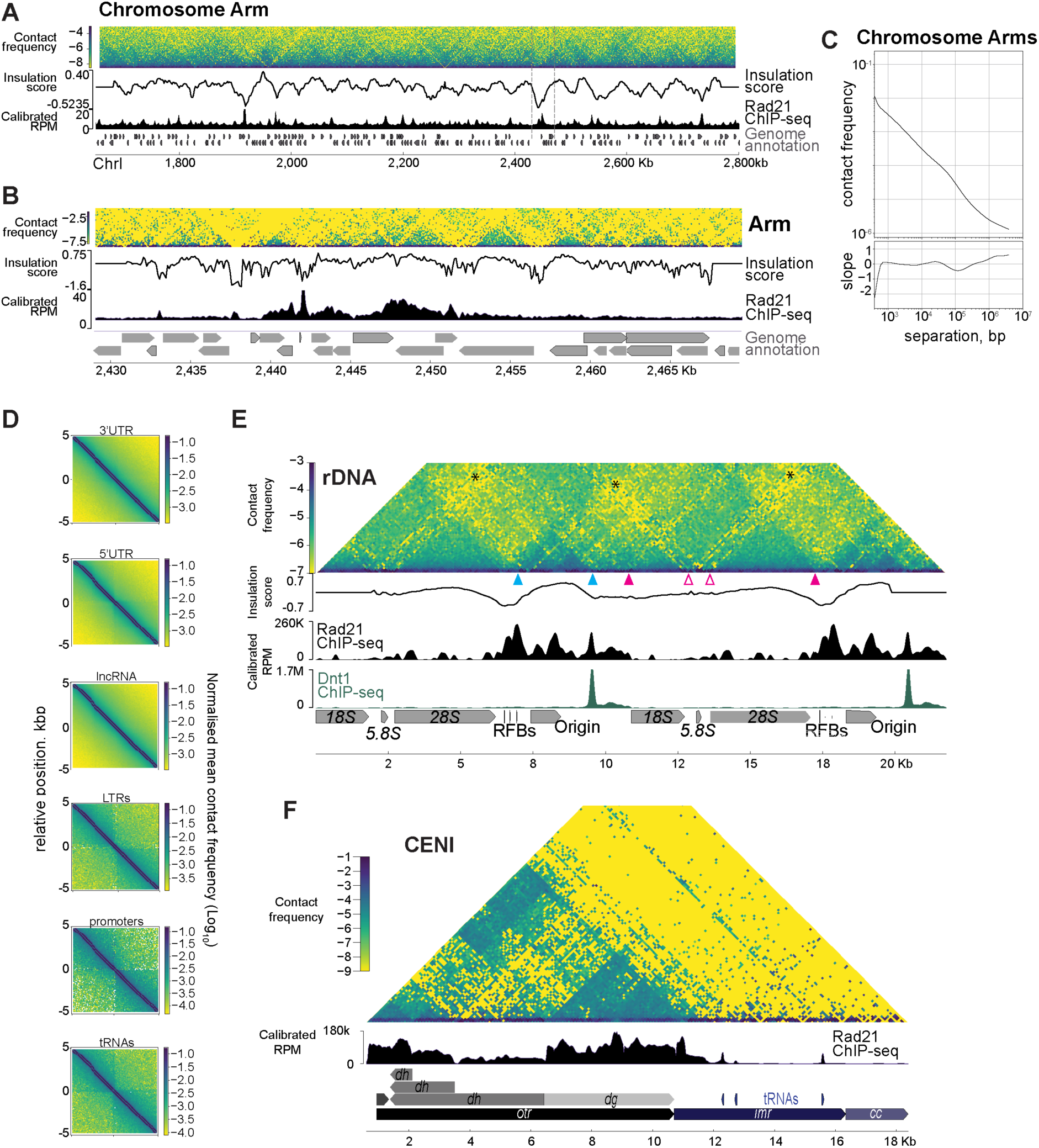
Cohesin defines local chromatin architecture in *S pombe.* (A-D) 5’UTRs, LTRs, convergent genes and tRNAs form boundaries of small locally interacting domains on chromosome arms. MicroC-XL matrix for wild type (AMfy29) showing a ∼1.2Mb region of chromosome I (A), grey lines denote the 135kb region shown in (B). Pileups of MicroC-XL data, centred on annotated genome features (C). Contact probability curve and its derivative plotted with pericentromeres and rDNA regions excluded (D). RFBs and Dnt1 define boundaries of rDNA domains in *S. pombe.* Contact matrix aligned to two complete repeats of the rDNA (E) Structure of centromeres in *S. pombe*. Arrowheads represent regions of interest referred to in the text. (F) contact matrix reads aggregated on the left side of Centromere I; different lengths of *dh* repeat from other centromeres are denoted,. See also Fig. S4.

To visualise rDNA chromatin interactions, we mapped MicroC-XL contacts to a custom reference genome containing two tandem rDNA repeats derived from the complete repeat on the right side of chromosome III. As the rDNA is highly repetitive, interactions represent aggregated multi-mapping reads, generating an rDNA-normalised contact matrix that accounts for potential differences in rDNA copy number between libraries. In wild type, we observe a strong domain of interactions spanning across most of the NTS region, which is enriched with cohesin and Dnt1 (Fig. 4E). Boundaries of this domain are positioned at the leftmost RFB (*RFB3*) and the strongest Dnt1 peak and coincide with two strong peaks of cohesin (Fig. 4E; cyan arrowheads). A larger, less prominent, domain spans across the rDNA genes and two ITS regions, which is also subdivided into gene-specific domains (Fig. 4E; filled and open magenta arrowheads, respectively). Importantly, despite the high condensation of the rDNA, contacts between the NTS and ITS domains are not enriched, indicating that the rDNA genes and cohesin-rich NTS regions occupy distinct spatial compartments within at least the majority of repeats (Fig. 4E, asterisks).

We used a similar approach to observe interactions at centromeres, assembling a custom genome file containing only the left half of centromere I onto which we mapped genomic reads in aggregate. The heterogeneity in *dh* repeat length across centromeres is reflected in the pattern of cohesin occupancy and MicroC-XL contacts (Fig. 4F). Nevertheless, we observed prominent domains of interaction within the outer *dg* and *dh* repeats where cohesin is concentrated, while the cohesin-depleted core centromere lacked extensive interaction domains (Fig. 4F). Within the outer repeats, stripes of interaction tended to emanate from cohesin peaks. Notably, cohesin peaks at tRNA genes in the inner repeats exhibited diagonal stripes extending away from the core centromere (Fig. 4F). Although the repetitive nature of these regions precludes definitive conclusions about their detailed organisation, our analysis indicates that cohesin is enriched at domain borders in rDNA and pericentromeres.

### Chromosome arm structure is largely robust to loss of cohesin targeting

Next, we asked how loss of targeted cohesin loading alters chromosome structure. We generated MicroC-XL maps for *ssl3-1E2A* cells, which lose cohesin targeting; and *mis4-1K1F* cells, where cohesin loading is independent of Ssl3 so that at least a fraction of cohesin is also predicted to be untargeted. We also generated maps for the *ssl3-1E2A mis4-1K1F* double mutant, which combines loss of cohesin targeting with the increased cohesin loading capacity conferred by *mis4-1K1F.* At the whole-chromosome level, we observed no major changes in overall chromosome organisation compared to wild type (Fig. S5A,B). This is consistent with the modest effects on cohesin distribution along chromosome arms in these mutants (Fig. 2E). However, both *mis4-1K1F* and *mis4-1K1F ssl3-1E1A* cells displayed stronger short-range interactions, evident from the increased intensity and definition of contact map domains and higher contact frequencies over the 100-1000bp range in contact probability curves generated after excluding the rDNA and pericentromeric regions (Fig. S5B,C). In contrast, contact probabilities curves for chromosomal arms were essentially identical between wild type and *ssl3-1E2A* (Fig. S5C). Therefore, cohesin targeting via the Ssl3 patch is largely dispensable for chromosome arm structure, whereas Ssl3-bypass mutant Mis4-1K1F strengthens short-range chromosome contacts without substantially altering steady-state cohesin distribution. Notably, *mis4-1K1F* does alter the genomic distribution of Mis4 (Fig. 2H), but not Rad21 (Fig. 2E) suggesting that changes in cohesin loading position or dynamics, rather than its final chromosomal distribution, underlie the altered chromosome architecture.

### Cohesin targeting organises pericentromeres

While cohesin targeting via the Ssl3 patch is largely dispensable for cohesin patterning on chromosome arms, cohesin loading at specialised domains, the pericentromeres and rDNA, was largely abolished in *ssl3-1E1A* cells (Fig. 1G-I). Accordingly, we anticipated that the structural organisation of specialised domains would be altered in these cells. To examine the effect on pericentromere structure, we mapped MicroC-XL data onto a custom genome comprising the left side of the chromosome I regional centromere. Repetitive sequences, both within and between pericentromeres, preclude interpretation of these MicroC-XL maps as representing the native structure of an individual pericentromere. Nevertheless, comparison of mutant and wild-type maps enables broad conclusions about how loss of cohesin targeting affects the structural organisation of this region. In *ssl3-1E2A* cells, the interaction pattern became more diffuse (Fig. 5A), with a reduction in very short-range interactions evident from the enrichment of wild type (blue) contacts along the base of the difference map (Fig. 5B). Concomitantly, contacts over slightly longer genomic distances increased throughout the pericentromere, producing an overall enrichment of contacts in *ssl3-1EA* (red) in the difference map (Fig. 5B). These changes were accompanied by increased interactions between the *dh* and *dg* outer repeats, consistent with a loss of focused local interactions (Fig. 5A). To examine these changes in more detail, we performed Virtual 4C using viewpoints within the outer repeats, inner repeats or core centromere (Fig. 5C-E). This confirmed that contacts originating from both the outer and inner repeats became broader and more dispersed throughout these regions, with some extending into adjacent domains (Fig. 5C,D). Similarly, interaction peaks at the core centromere were diminished in the *ssl3-1E2A* cells, suggesting a loss of precise loop positioning (Fig. 5E). We also noticed that interaction stripes emanating from tRNAs within the inner repeats became more prominent in *ssl3-1E2A* cells, with a new stripe emerging from the second leftmost tRNA through the core centromere (Fig. 5A; magenta arrow). These observations suggest that Ssl3-mediated positioning insulates tRNA interactions and restricts contacts between the inner repeats and central core.

**Figure 5.**
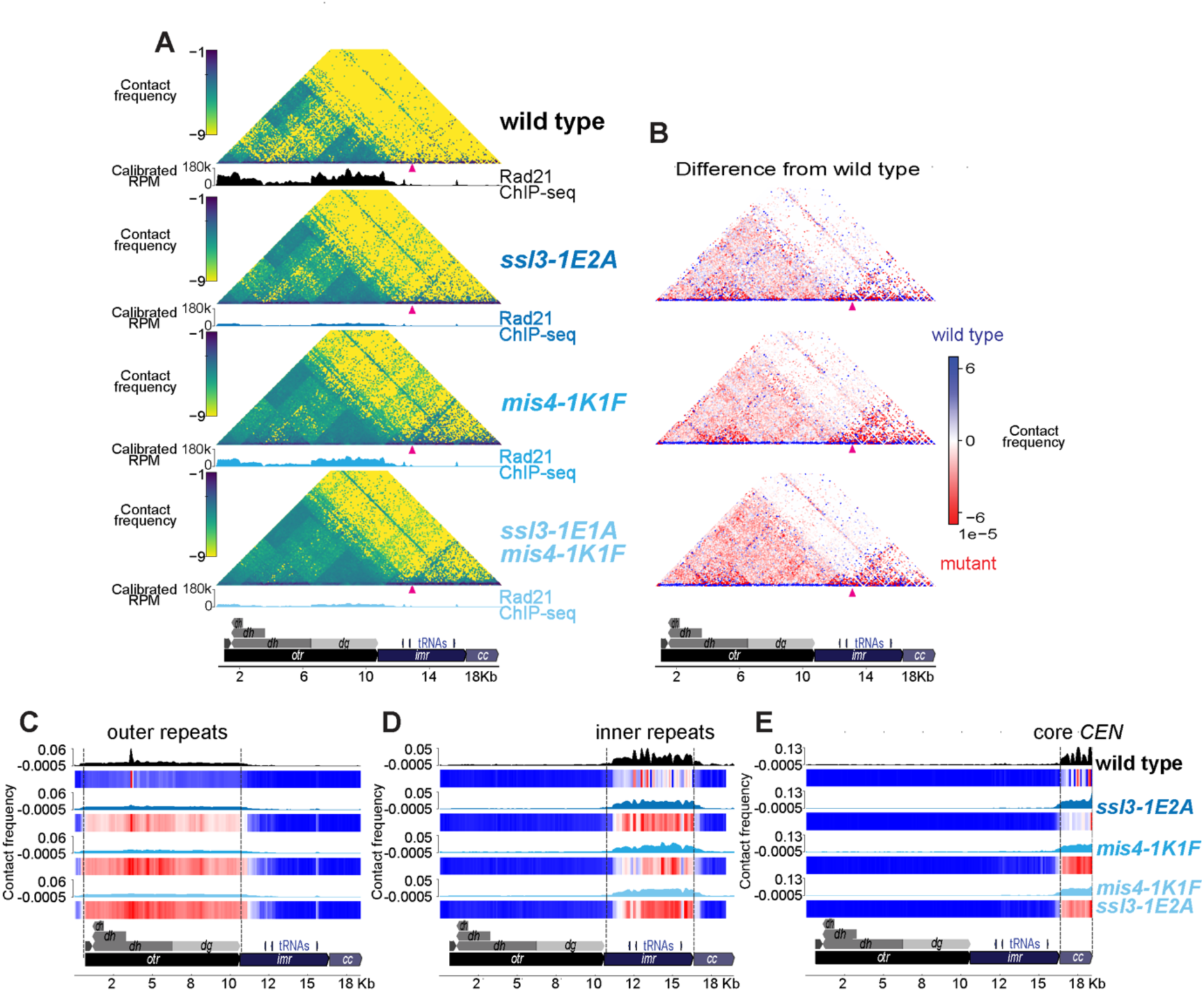
Relaxing spatial constraints on cohesin loading compromises pericentromere structure. Contact matrices with ChIP-seq profiles showing Rad21 enrichment (A). Difference plots show enriched contacts in wild type (blue) versus mutants (red) (C). (C-E) Virtual 4C of the contact matrices in A. Viewpoints (grey lines) encompass the outer repeats (C), inner repeats (D) and core centromere (E). Data is shown as filled line plots of interaction frequency (top panels) and heatmaps (bottom panels) denoting the proportion (highest red; lowest blue)) of contacts at 300bp resolution across the left side of CEN I. See also Fig S5.

Interestingly, the effects of *mis4-1K1F* and *mis4-1K1F ssl3-1E2A* on pericentromere organisation closely resembled those observed in *ssl3-1E2A* cells, but were substantially more pronounced (Fig. 5A-E). Focused short-range interactions were further diminished, contacts between the *dg* and *dh* outer repeats were strengthened, and interaction stripes emanating from the tRNAs became more prominent. These phenotypes were strongest in the double mutant, suggesting that complete loss of Ssl3-dependent cohesin targeting leads to further loss of positional specification in *mis4-1K1F* cells. In addition, and unlike *ssl3-1E2A* alone, *mis4-1K1F* markedly increased interactions throughout the core centromere, both in the presence and absence of *ssl3-1E2A* (Fig. 5A,B,E). Rather than forming discrete interaction peaks, these contacts were broadly distributed across the core centromere (Fig. 5E), indicating that the *mis4-1K1F* bypass allele promotes cohesin-dependent chromosome folding but with reduced spatial precision.

### Targeted cohesin loading defines domain-specific chromosome architecture at the rDNA

Next, we sought to understand how cohesin targeting shapes rDNA organisation. Mapping wild-type MicroC-XL onto two consecutive rDNA repeats identified discrete interaction domains within the transcription unit and NTS rDNA regions, with the Dnt1 peak and RFBs demarcating boundaries between them (Fig. 4E). To place cohesin-dependent organisation in the context of other architectural features, we additionally performed ChIP-Seq of the condensin subunit Cnd2, which is known to play important roles in rDNA structure and function. Cnd2 exhibited a localisation profile broadly similar to cohesin across the rDNA, but with distinct differences in enrichment. The strongest Cnd2 signal occurred over the replication origin, enrichment at the Dnt1 peak was weaker than for cohesin, and occupancy across the three RFBs was more uniform (Fig. 6A).

**Figure 6.**
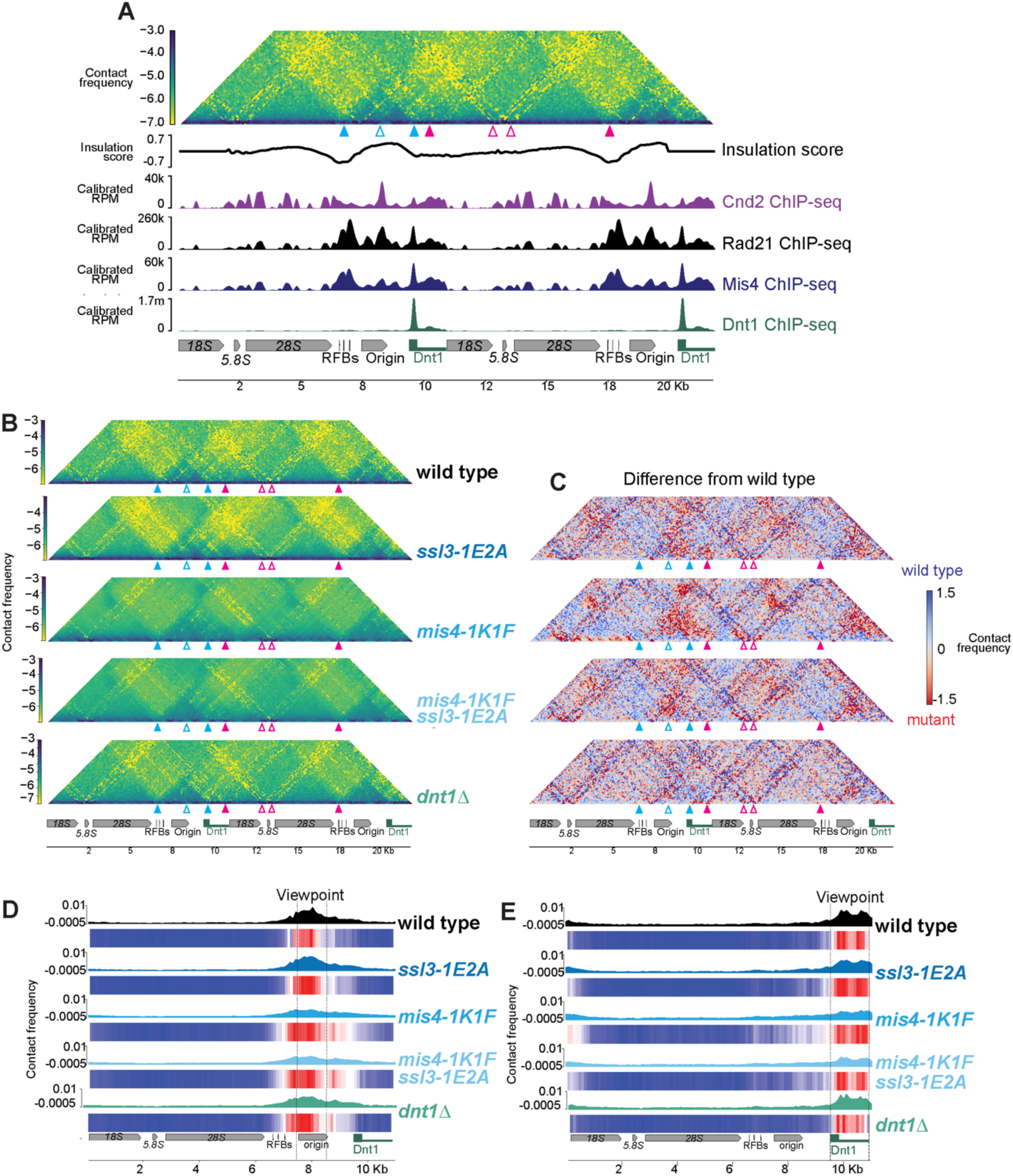
Relaxed cohesin positioning significantly alters rDNA architecture. Cohesin, Mis4 and condensin are enriched at rDNA (A). Wild type (AMfy29) contact matrix of rDNA with ChIP-seq profiles showing enrichment of condensin (AMfy1862, *CND2-6HA*), cohesin (AMfy638; *RAD21-6HA*), Mis4 (AMfy659, *MIS4-6HA*) and Dnt1 (AMfy2203, *DNT1-9MYC*). MicroC-XL contact matrices (B) and difference plots (C) showing enriched contacts in wild type (blue) and mutants (red) normalised to total plotted sequencing reads for each matrix. Virtual 4C of the contact matrices in A with viewpoints (grey lines) at the replication origin (D) and Dnt1 rich region (E). Filled line plots depict interaction frequency (top panels) and heatmaps (bottom panels) denote the proportion (highest red; lowest blue) of contacts across a single rDNA repeat at 300bp resolution. See also Figure S6.

Loss of the Ssl3 patch resulted in clear alterations to rDNA organisation (Fig. 6B-E). Although the two major domains encompassing the NTS and transcription unit were retained, interactions between them were reduced compared with wild type. Instead, interaction stripes emanating from the RFBs (leftmost filled cyan arrowhead) became more prominent, resulting in stronger insulation between rDNA repeats. There were also more local effects within the NTS, with stripes emerging from the replication origin (open cyan arrowhead) also increasing in strength, extending across the Dnt1-binding region. Because Cnd2 is enriched at these same sites (Fig. 6A), these observations suggest that condensin establishes the major domain boundaries within the rDNA, whereas cohesin targeted by the Ssl3 patch normally confines condensin-associated interactions to defined chromosomal intervals. In the absence of cohesin, these stripes extend further along the chromosome while forming stronger boundaries that reduce contacts between neighbouring domains.

The Ssl3-bypass mutant *mis4-1K1F*, as well as the *ssl3-1E2A mis4-1K1F* double mutant, also exhibited stronger RFB- and origin-associated interaction stripes extending across the Dnt1-binding region (Fig. 6B,C; cyan filled and unfilled arrowheads, respectively). The spreading of contacts was particularly evident in Virtual 4C using either the replication origin or Dnt1-binding region as viewpoints (Fig. 6D,E). This mirrors the effects of *mis4-1K1F* on pericentromere organisation and suggests that bypass of Ssl3 enhances cohesin-mediated chromosome folding while reducing its spatial constraint. Consistent with this, boundaries between the transcription unit and NTS were weakened in both mutants, resulting in more frequent contacts between these domains (Fig. 6B,C).

Our data support a model in which cohesin enrichment within the rDNA is mediated, at least in part, by the interaction between Dnt1 and the conserved Ssl3 patch. Dnt1 forms a prominent peak between consecutive rDNA repeats, which coincides with a domain boundary in wild type cells (Fig. 6A, rightmost cyan arrowhead). In *ssl3-1E2A* cells, this boundary was overcome unidirectionally with RFB-associated interaction stripes extending into the NTS and accompanied by reduced contacts between rDNA domains (Fig. 6B). In contrast, interactions between the transcription unit and NTS were modestly increased in *dnt1Δ* (Fig. 6B,C,E), suggesting that Dnt1 helps to insulate rDNA repeats in a manner independent of cohesin targeting. However, like *ssl3-1E2A*, *dnt1Δ* cells showed a small loss of interactions in the NTS region where Dnt1 is normally enriched (Fig. 6B-E; open cyan triangle and rightmost filled cyan triangle). This was accompanied by a modest increase of non-specific contacts across the rDNA region, consistent with redistribution of Ssl3/Mis4-dependent cohesin loading away from Dnt1 sites (Fig. 6B-E). Similarly, although centromere and arm architecture are not detectably altered in *dnt1Δ*, the density of centromere contacts is slightly increased, likely reflecting redistribution of cohesin to centromeres in this mutant (Fig. S6D).

Together, these findings indicate that Dnt1 and the Ssl3 patch spatially regulate cohesin-dependent chromosome folding at the rDNA. Conversely, bypass of Ssl3 by *mis4-1K1F* relaxes these spatial constraints, allowing cohesin-dependent chromosome interactions to extend beyond their normal boundaries. Thus, targeted regulation of the Mis4/Ssl3 loader acts not simply to enrich cohesin at specific loci, but to define where cohesin activity shapes chromosome architecture.

## Discussion

Here we show that spatial targeting of cohesin loading is a key determinant of chromosome architecture. By separating activation of the cohesin loader Mis4 from its positioning by Ssl3, we demonstrate that targeted loading at specialised chromosomal domains is required to focus cohesin-mediated interactions and establish local chromosome organisation. We further identify the nucleolar protein Dnt1 as a receptor that recruits the Mis4-Ssl3 complex to the rDNA locus, providing a mechanistic example of how chromosomal features shape local genome structure.

### Targeted loading defines specialised chromosomal domains

Our analysis reveals that cohesin organisation differs markedly across chromosomal domains in *S. pombe*. At pericentromeres, Rad21 is enriched over heterochromatic outer repeats, whereas Mis4 is concentrated at the central core and inner repeats, indicating that distinct cohesin populations occupy specialised chromosomal environments. Disruption of kinetochore function primarily affected Mis4 localisation within the core centromere, whereas loss of heterochromatin reduced cohesin enrichment across the pericentromeric outer repeats, as reported (Bernard *et al*, 2001; Nonaka *et al*, 2002; Bailis *et al*, 2003). Together, these observations indicate that distinct chromosomal features independently regulate cohesin loading and retention within different regions of the pericentromere, while collectively establishing its overall organisation. In contrast, cohesin and Mis4 display highly overlapping localisation across the rDNA locus, with enrichment concentrated in the NTS region. This pattern is consistent with cohesin activity being focused at this region and accordingly, we provide evidence that targeted cohesin loading plays important roles in local domain interactions within this region, as well as limiting its insulation from the adjacent transcribed region. More generally, these observations indicate that cohesin localisation is shaped by domain-specific recruitment mechanisms and that targeted loading represents a key organising principle underlying specialised chromosomal domains.

### A conserved Ssl3 interface positions cohesin loader activity

Our findings reveal that the conserved Scc4/Ssl3 interface functions as a targeting module that positions the cohesin loader at specific chromosomal sites. Mutation of this conserved surface abolishes cohesin enrichment at centromeres and rDNA while leaving most chromosome arm cohesin largely intact (Fig. 1D-F), indicating that this interface is specifically required for targeted cohesin loading rather than general cohesin association with chromosomes. Analysis of the Mis4 bypass mutant allowed us to separate two functions of the cohesin loader: activation of cohesin loading and spatial targeting of that activity. The *mis4-1K1F* allele restores cohesin loading in the absence of Ssl3-dependent activation but fails to restore cohesin enrichment at centromeres or rDNA when the conserved Ssl3 targeting surface is disrupted (Fig. 2A,B). Moreover, although mis4-1K1F alters Mis4 localisation, steady-state cohesin occupancy changes comparatively little, consistent with the idea that cohesin redistributes after loading to preferred chromosomal positions. These observations indicate that chromosome architecture depends less on where cohesin ultimately accumulates than on where loader activity is initiated.

Our identification of the nucleolar protein Dnt1 as a receptor for the Mis4–Ssl3 complex provides a mechanistic example of this targeting principle. Dnt1 localises to a discrete site within the rDNA NTS and is required to focus Mis4 and cohesin recruitment at this locus. This function is analogous to the recruitment of Scc2-Scc4 by Ctf19^CENP-P^ at the *S. cerevisiae* kinetochore (Hinshaw *et al*, 2017), supporting the idea that diverse chromosomal receptors engage the conserved Ssl3 interface to spatially regulate cohesin loading. Whether DDK or other kinases regulate this Ssl3-receptor interaction is unclear. While DDK was reported to promote cohesin recruitment to pericentromeres (Bailis *et al*, 2003), we obtained no evidence that Dfp1(1-136), which lacks the Swi6 interaction motif, impaired cohesin enrichment in pericentromeres. Consistently, a later study reported no increase in lagging chromosomes (a marker of cohesion defects) in the *swi6-3A* mutant which similarly abolishes interaction with Dpf1 (Shen & Forsburg, 2019). Therefore, DDK may promote pericentromeric cohesin enrichment independent of its interaction with Swi6 and future work will be required to understand the role of DDK in cohesin targeting in *S. pombe*.

Overall, these findings support a model in which the conserved Scc4/Ssl3 interface serves as a modular docking surface that engages diverse chromosomal receptors to direct cohesin loading to specialised genomic regions. Our findings indicate the existence of additional, as-yet-to be identified, receptors, at least in the kinetochore and pericentromeric heterochromatin that target cohesin to these locations. Identification of these receptors will be an important future priority. Notably, mutations affecting the corresponding surface in human MAU2 are associated with the developmental disorder Cornelia de Lange syndrome (Parenti *et al*, 2020, 2025), suggesting that impairment of targeted cohesin loading may also contribute to pathology.

### Spatial control of cohesin loading defines chromosomal domains

Our MicroC-XL analyses demonstrate that spatial targeting of cohesin loading plays a key role in defining chromosome architecture. Loss of the Ssl3 targeting surface did not substantially alter global chromosome arm organisation but produced pronounced changes in specialised domains where cohesin loading is normally targeted, namely pericentromeres and rDNA. In these regions, chromosome interactions became less spatially constrained, despite the persistence of large-scale domain organisation. These observations indicate that in *S. pombe,* targeted cohesin loading primarily organises interactions within chromosomal domains rather than establishing the domains themselves. The *mis4-1K1F* bypass mutant provides further mechanistic insight. Although cohesin occupancy changed relatively little, chromosome interactions became broader and less focused, indicating that bypass of Ssl3 relaxes the positional control normally imposed. We propose that when cohesin loading is no longer spatially restricted, cohesin-dependent chromosome folding is initiated over a broader range of chromosomal sites, reducing the positional reproducibility of chromosome interactions observed across a cell population, or, where appropriate, between individual repeats within the same cell.

Together, our findings establish spatial regulation of cohesin loader activity as a fundamental mechanism for generating chromosome domain diversity. Rather than simply determining where cohesin accumulates, chromosomal receptors acting through the conserved Ssl3 interface define where cohesin becomes competent to shape chromosome architecture. This provides a general framework for understanding how distinct chromosomal domains acquire specialised three-dimensional organisation despite sharing the same conserved cohesin machinery. Identifying additional receptors that engage this conserved loader interface will be important for understanding how the fine-scale genome organisation is specified across eukaryotes.

## Acknowledgements

We are grateful to Lori Koch and Tania Auchynnikava for help with proteomics analysis and to Angelika Amon, Kevin Hardwick, Alison Pidoux, David Tollervey and Robin Allshire for *S. pombe* strains and plasmids. We thank Robin Allshire, Stephen Hinshaw, Alison Pidoux, Jean-Paul Javerzat and members of the Marston group for helpful discussions. We gratefully acknowledge the Wellcome Discovery Research Platform for Hidden Cell Biology Proteomics and Bioinformatics Cores. This work was funded through Wellcome Investigator [220780] and Discovery Awards to AM [319314], a BBSRC EASTBIO studentship to MPJ, core funding for the Wellcome Centre for Cell Biology [203149] and a Wellcome Discovery Research Platform Award [226791].

## Author contributions

Conceptualization – HR, MPJ and AM; Data curation – HR, MPJ, DR, SW and CS; Funding Acquisition – AM; Formal Analysis – HR, MPJ, AM, CS; Investigation – HR, MPJ, DR, SW and CS; Software – DR, SW; Supervision – AM; Visualization – HR, AM; Writing – original draft – HR, AM; Writing – review and editing – HR and AM with input from all authors

## Methods

### Yeast strains, plasmids and culture conditions

*S. pombe* and calibration *Saccharomyces cerevisiae* strains used in this study are given in Table 1.

Cycling cells were routinely grown in 1X YES at 32°C with shaking. For IP-Mass spectrometry cells were grown to OD_600_=1.5-6 in 4X YES to support logarithmic growth to high density. For experiments using temperature sensitive strains, cultures were grown to mid-log phase at 25°C before being shifted to 36°C for 4 (*ssl3-29*) or 6 hours (*fta2-291*).

Gene deletion and tagging was performed by standard PCR and transformation-based methods. Mutations were introduced by transforming with a DNA fragment amplified from a plasmid containing the full-length gene mutated using the Quickchange XL kit (Agilent Technologies 200517).

### Calibrated ChIP-seq

Calibrated ChIP-sequencing was performed as previously described (Paldi *et al*, 2020; Barton *et al*, 2022) with the following modifications. Cycling cultures of *S. pombe* were crosslinked for 2 hours with 1% formaldehyde as described previously. For calibration, a culture of *S. cerevisiae* with a protein tagged by the same epitope (Table1) was grown to OD_600_=0.25-0.3, frozen as described, and spiked into the *S. pombe* sample at a ratio of ∼2 volumes *S pombe*: 1 *S. cerevisiae*. Samples were lysed by 3 cycles of 30 seconds on high using FastPrep24™ Classic in the presence of silica beads and 0.5% SDS. Crosslinked DNA was sheared to an average size of 200-500bp by two rounds of 30 cycles 30 seconds sonication on high on Diagenode Bioruptor Plus bath sonicator temperature controlled to 4°C by a Diagenode Minichiller 300. Immunoprecipitation was performed as described with 12CA5 (Roche anti-HA, 11666606001) for Rad21 and Mis4 or anti-Myc antibody (9E10 anti-Myc Antibodies.com A278887).

At least 12.5M reads per immunoprecipitation and 2.5M reads per Input control was obtained by sequencing using llumina MiniSeq with MiniSeq High output reagent kit (150-cycles) (Illumina FC-420-1002) or by submitting pre-pooled libraries through Genewiz Azenta NGS services using Illumina NovoSeq X Plus platform.

Read mapping, normalisation and data visualisation was performed as previously described (Barton *et al*, 2022). Pileups and peak calling were performed using deeptools 3.5.3; detailed scripts are available at https://github.com/hollierowlands/Rowlands2026.

### Co-immunoprecipitation-Mass spectrometry

Cycling cultures of *S. pombe* were grown to OD_595_1.5-6 in 4X YES to support logarithmic growth at high cellular density. Cells were washed twice in H0.15 buffer (25mM HEPES, 2mM MgCl_2_ 0.1mM EDTA, 0.5mM EGTA, 15% Glycerol, 0.1% Nonidet-P40, 150mM KCl) resuspended at 2:1 weight/volume H0.15 plus inhibitors (2mM β-glycerophosphate, 1mM sodium pyrophosphate, 5mM sodium fluoride, 100μM sodium orthovanadate, 2mM AEBSF, 0.2μM microcystin, 1mM NEM, 1X Roche cOmplete, EDTA-free protease inhibitor cocktail).

Cell pellets were ground to a fine powder using SPEX Freezer Mill 6870 precooled for 2 minutes, then run at 10 cycles per second for a total of 8 cycles, 2 minutes ON/2 minutes OFF. Lysates were stored at -80°C until immunoprecipitation.

Merck Rabbit IgG (12370) were pre-conjugated to Protein G Dynabeads by incubating in a borate buffer (40.4mM boric acid, 40.1mM Sodium tetraborate decahydrate) containing 20mM DMP for 30 minutes at 4°C, before being washed into mild lysis buffer (50mM Tris pH 8, 150mM KCl, 0.1% TritonX-100 Prior to immunoprecipitation beads were washed twice into IP buffer. For immunoprecipitation, 1g of lysate was resuspended in 2mL of IP buffer plus inhibitors and incubated on ice with periodic rotation until completely defrosted. Cell debris were cleared by centrifugation and the supernatant was taken for immunoprecipitation with 1mg/g IgG-conjugated beads (Ssl3) or 0.5mg/g for Mis4. Immunoprecipitation reactions were carried out at 4°C for 1.5 hours with rotation, then beads were washed 3 times with IP buffer (Ssl3) or 4 times in PBS for Mis4. After the final wash, IPed proteins were eluted using 0.1% Rapigest (TE pH 8) solution by heating for 15 minutes at 60°C.

Immunoprecipitated proteins were prepared for mass spectrometry using the filter-aided sample preparation (FASP) method (Wiśniewski *et al*, 2009) with minor modifications. Proteins were reduced with dithiothreitol (DTT), alkylated with iodoacetamide, and processed on FASP filters (Sartorius, VN01H21). Following sequential washes with urea buffer and 50 mM ammonium bicarbonate (ABC), proteins were digested on-filter overnight at 37°C using sequencing-grade modified trypsin (Pierce). Peptides were recovered by centrifugation, washed with 50 mM ABC, and acidified with trifluoroacetic acid (TFA). Peptides were desalted using homemade C18 StageTips (Rappsilber *et al*, 2003) prepared from three Empore C18 extraction discs. StageTips were conditioned with methanol, 80% acetonitrile/0.1% TFA, and 0.1% TFA before loading acidified peptide samples (pH 1–2). Following washing with 0.1% TFA, StageTips were stored at −20°C until analysis. Peptides were eluted from the StageTips and analysed by nanoLC–MS/MS on an Orbitrap Fusion Lumos Tribrid mass spectrometer (Thermo Fisher Scientific), as described by (Blyth *et al*, 2018) or on Orbitrap Exploris™ 480 (Thermo Fisher Scientific, UK) on a Data Independent Acquisition (DIA) mode, coupled on-line, to an Ultimate 3000 HPLC (Dionex, Thermo Fisher Scientific, UK), as described by (Koch *et al*, 2026, 2024).

Detailed analysis scripts for the Mis4-SZZ experiment can be found at https://github.com/lori-koch/Rowlands_2026.

For the experiment comparing Ssl3 interactors in wildtype and the Ssl3 patch mutant, samples were analysed using FragPipe v24 LFQ analysis to compare intensities between the samples, with variance stabilising normalisation, Perseus-type imputation and Benjamin-Hochberg FDR correction.

### MicroC-XL

300mL cultures of cycling cells were grown to OD_595_0.55 and fixed for 15 minutes in 3% formaldehyde at 30°C with gentle rotation. Formaldehyde was quenched by addition of 125mM glycine. Fixed cells were isolated by centrifugation at 4000rpm for 4 minutes at 4°C, and washed once with cold water before being resuspended in Buffer Z (1M sorbitol, 50 mM Tris-HCl pH 7.4, 10 mM β-mercaptoethanol) and incubated at 30°C with gentle rotation in the presence of 417mg/mL Zymolyase-100T (AMS Biotechnology Ltd 120493-1) until completely digested. Spheroplasts were washed once in PBS, then fixed with 300μM Disuccinyl glucinate (ThermoFisher Scientific H58208.MC) long crosslinker in PBS for 40 minutes at 30°C with gentle rotation before being quenched by the addition of 125mM glycine at room temperature. Fixed spheroplasts were washed in PBS before being snap frozen and stored at -70°C until MNase digestion.

Fixed spheroplasts were resuspended in Buffer MB1 (50mM NaCl, 10mM Tris, 5mM MgCl2, 1mM CaCl2, 0.5mM Spermidine, 1.43mM B-Me, 0.005% NP-40) and incubated on ice for 20 minutes before being subjected to MNase (Worthington Biochemical LS004798) titration. After treatment with MNase for 20 minutes at 37°C, enzymatic activity was stopped by the addition of 2.5mM EGTA and incubating at 65°C for 10 minutes. Digested pellets were washed twice in buffer MB2 (50 mM NaCl, 10 mM Tris-HCl pH 7.5, 10 mM MgCl2) and 1:250 of sample was taken and reverse crosslinked in the presence of 200μg/mL RNase A (Fisher EN0531), 2mg/mL Proteinase K, 0.5% SDS, 1XTE and incubated at 55°C for 15 minutes, then 65°C overnight. Samples were purified using Promega Wizard PCR cleanup kit (Promega A9282) and run on a 2% agarose gel in 1X TAE to check quality of MNase digestion. A sample with ∼90% monosomes/10% disomes was selected for proximity ligation.

In preparation for proximity ligation, DNA ends were subjected to treatment with T4 Polynucleotide Kinase (1X NEBuffer r2.1, 2mM ATP, 3mM DTT, 25U NEB T4 PNK(M0201L)) for 15 minutes at 37°C. Followed by a further 15 minutes incubation in the presence of 25U NEB Klenow (Large) Fragment (NEB M0210L). End labelling was performed by adding 67μM biotin-14-dATP (Jena Bioscience NU-835-BIO14-S), biotin-11-dCTP (B8346-APE-50uL), 67μM each dTTP and dGTP (NEB N0446S), 1X NEB T4 DNA ligase buffer and 33.3μg/mL BSA (NEB B9200S) and incubating for 45 minutes at 25°C. The end labelling reaction was stopped by the addition of 40μM EDTA and heat inactivation at 65°C for 20 minutes. Pellets were washed once in MB3 (50mM Tris pH 7.5, 10mM MgCl_2_) before proceeding to proximity ligation. Proximity ligation was performed at room temperature for 4 hours in the presence of 1X T4 DNA ligase buffer (NEB M202L), 100μg/mL BSA, 5000U T4 DNA ligase (NEB M0202L). Unligated biotinylated DNA ends were removed by Exonuclease III (NEB M0206L) incubated at 37°C for 15 minutes.

Crosslinks were reversed by adding 1X TE, 400ug/mL Proteinase K (VWR EO0492), 40μg/mL RNase A (Fisher EN0531), 1% SDS and incubating at 55°C for 15 minutes then 65°C overnight with agitation. DNA was isolated first by performing an extraction with phenol/chloroform/isoamyl alcohol and then purifying the top aqueous phase through a ZymoClean DNA Clean and Concentrator (Zymo Research D4014) kit. The purified DNA was run on a 3.5% Nuseive GTG Agarose (Lonza 50084) gel in 1X TAE, from which the band containing ligated disomes was isolated and purified using ZymoClean Gel DNA Recovery kit (Zymo Research D4001). The entirety of the isolated DNA was used to prepare MicroC-XL libraries

End repair was performed using Qiagen End Repair Kit (Qiagen Y9140-CL-L) or End-It End Repair Kit (Biosearch Technologies ER81050). For the Qiagen End Repair kit, reactions (1X End Repair buffer, 1X dNTP, 1uL Enzyme mix) were incubated at 20°C for 30 minutes, then 70°C for 10 minutes to inactivate enzymes. For End-It DNA End repair reactions (1X End-It Buffer, 1X ATP, 1X dNTP, 0.5μL End-It enzyme mix) were incubated for 45 minutes at 25°C then 70°C for 10 minutes. Biotinylated, ligated disomes were bound to Dynabeads^™^ MyOne C1 Streptavidin beads (ThermoFisher 65002) in 1X BW buffer (2.5 mM Tris-HCl pH 7.5, 25μM EDTA, 500mM NaCl) by incubation for 20 minutes at room temperature with rotation. Beads were washed twice in TBW buffer (0.1% Tween20, 5 mM Tris-HCl pH 7.5, 0.5 mM EDTA, 1 M NaCl) at 55°C for 5 minutes, then once with 10mM Tris pH 7.5.

Libraries were prepared from DNA on beads using NEB Next Ultra II Library Prep kit for Illumina (NEB 7645S) with NEBNext^®^ Multiplex Oligos for Illumina (NEB 7335S). Reactions were performed according to manufacturer’s instructions except that they were incubated with 1000rpm with interval shaking at 1000rpm with 15s ON/3 minutes OFF to prevent sedimentation of the beads. Folllowing adaptor ligation, unligated adaptors were removed by washing twice with 1X TBW then once with 10mM Tris pH 7.5. Library PCR was performed on-beads using PCR Biosystems 2X VeriFi^™^ Library Amplification Mix (PCR Biosystems PB72.10-05) according to manufacturer’s instructions with 8-12 cycles in a 50-100μL reaction and then library DNA was purified by two rounds of purification with 0.9X Beckman Coulter AMPure XP beads (Beckman Coulter A63881).

### Analysis of MicroC-XL data

Libraries were sequenced to produce at least 100M reads per sample by Novogene NGS pre-made library sequencing or Azenta Genewiz custom library sequencing platforms. Read alignment, filtering and quality control were performed using code from Dovetail Genomics (https://github.com/dovetail-genomics/Micro-C?tab=readme-ov-file). All code used in the analysis of MicroC-XL data are available at https://github.com/hollierowlands. Contact matrices were routinely made using Cooler v0.8.9 zoomify to produce a balanced cool file with multiple resolutions. For the rDNA, reads were mapped to a custom genome file containing two complete repeats from the right side of Chromosome III. Centromere matrices, and accompanying ChIP-seq profiles, were produced by mapping data to the left half of centromere I. The resulting matrices were balanced using Cooler v0.8.9 balance.

Visualisation of the matrices, virtual 4C and difference plots were created using Coolbox 0.3.9 except that HiGlass-Server v1.14.8 (Kerpedjiev et al. 2018) was used to generate a whole genome view. Cooltools 0.7.1 was used calculate and plot contact probability curves and their derivatives, and for the creation of pileups. Detailed analysis scripts can be found at https://github.com/hollierowlands/Rowlands2026.

---

**Figure S1.**
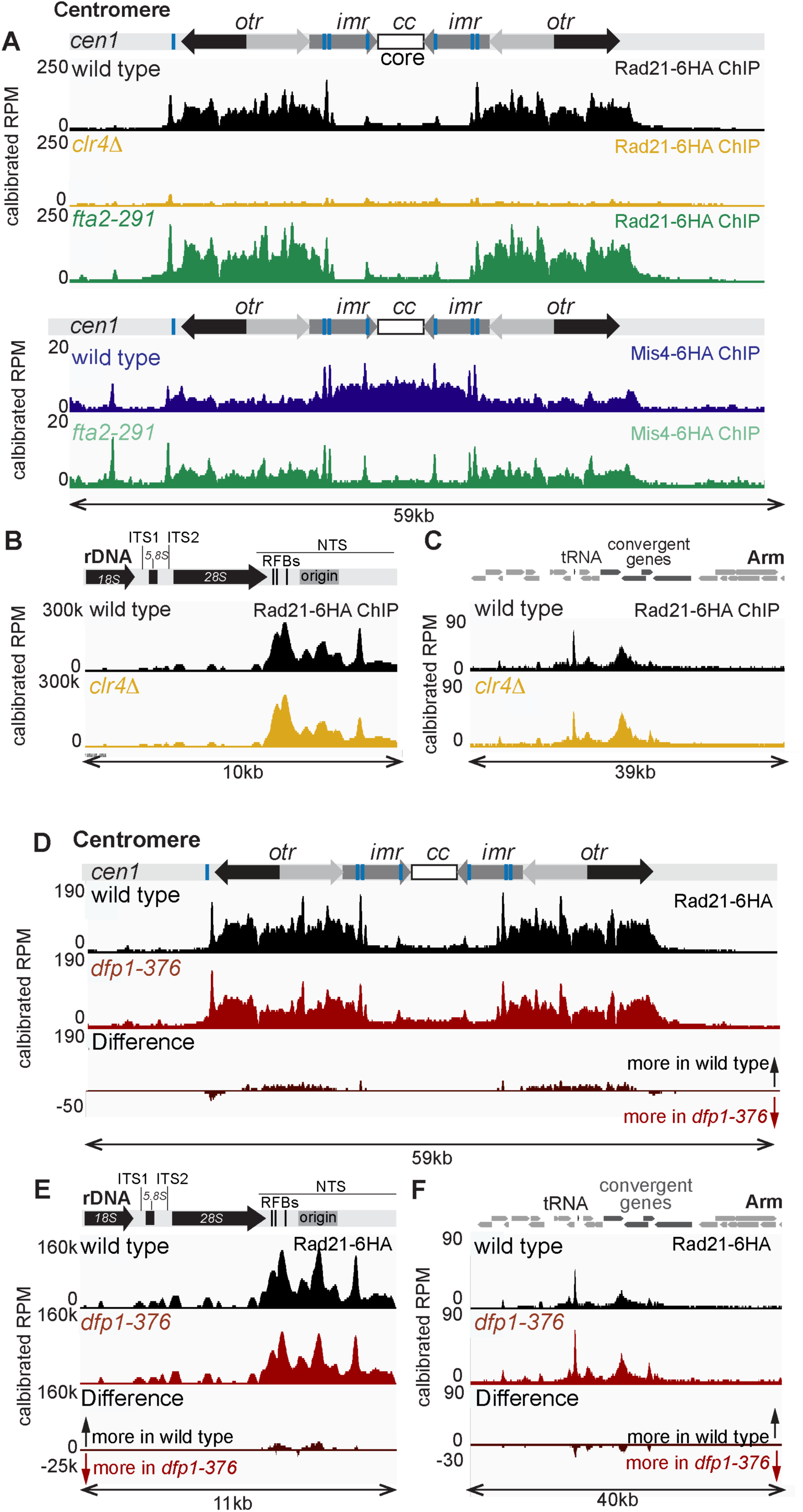
Importance of heterochromatin and kinetochores for cohesin and Mis4 targeting to chromosomal regions. (A-C) Effect of heterochromatin and kinetochores on cohesin and Mis4 localisation. Rad21-6HA or Mis4-6HA ChIP-Seq at pericentromeres (A), rDNA (B) or a chromosomal arm region (C) is shown for *clr4Δ* and/or temperature-sensitive *fta2-291* strains. Rad21-6HA strains were AMfy638 (wild type), AMfy1979 (*clr4Δ*) and AMfy2004 (*fta2-291*). Mis4-6HA strains were AMfy659 (wild type) and AMfy2260 (*fta2-291*). (D-F) Dfp1 C-terminal region is not required for cohesin enrichment in the pericentromeres (D), rDNA (E) but increases cohesin occupancy on arms (F). Strains were AMfy2107 (wild type) and AMfy2110 (*dfp1-(1-376)*.

**Figure S2.**
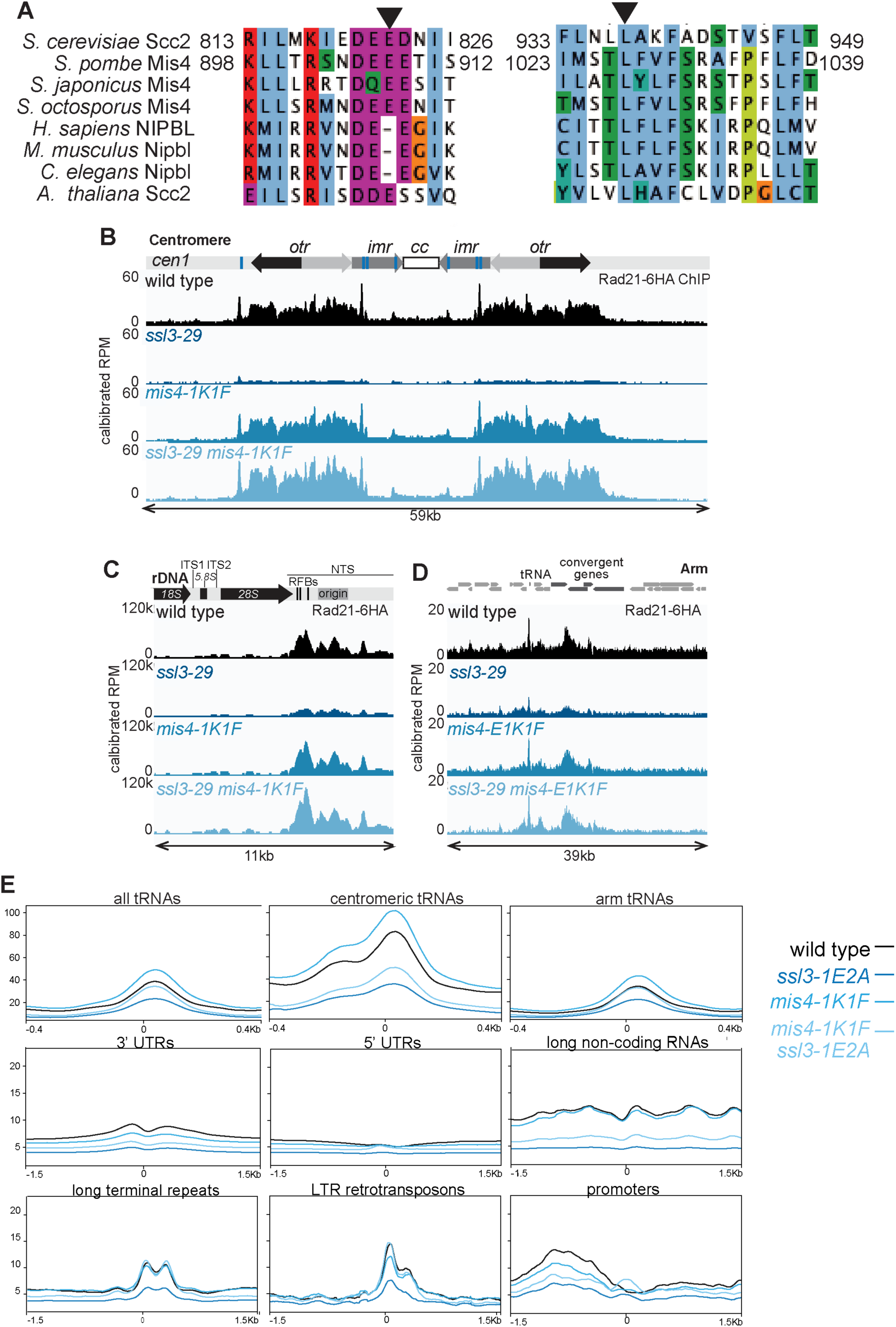
Hyperactivated Mis4 rescues cohesin localisation and lethality of the temperature-sensitive ssl3-29 allele. (A) Alignment of two regions of Scc2/Mis4/NIPBL proteins. Residues mutated in the hyperactive Scc2 and the corresponding residues in Mis4 are indicated by arrowheads. (B-D) Rad21-6HA ChIP-Seq at centromeres (B), rDNA (C) and a chromosomal arm region (D) showing wild type (AMfy2157), *ssl3-29* (AMfy2158), *mis4-E907K L1027F* (AMfy2163) and *ssl3-29 mis4-E907K L1027F* (AMfy2162). (E) *mis4-1K1F* rescues cohesin loss at tRNAs on chromosome arms but not specialised domains. ChIP-seq pileups of Rad21 occupancy centered on tRNAs, 3’ untranslated regions, 5’untranslated regions, long non-coding RNAs, long terminal repeats, LTR retrotransposons and promoters.

**Figure S3.**
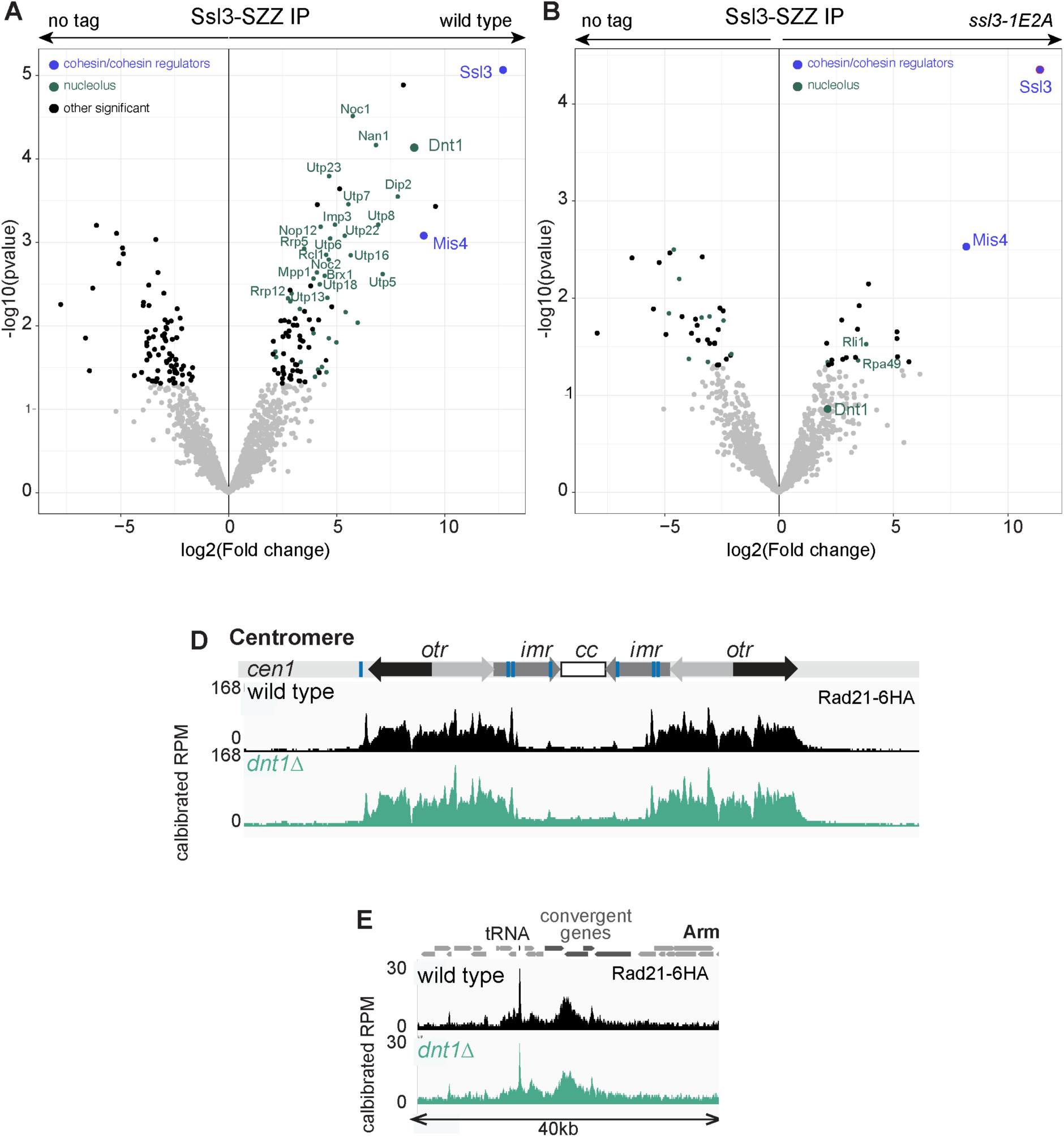
The conserved Ssl3 patch is required for interaction with nucleolar proteins but not Mis4. (A) Ssl3 interacts with Dnt1 and other nucleolar proteins. IP/mass spectrometry to identify Ssl3 (wild type) interactors. Strains AMfy205 (*ssl3-SZZ*) and AMfy29 (wild type no tag). Volcano plot compares proteins identified in Ssl3-SZZ immunoprecipitates compared to a no tag control. (B) Mutation of the Ssl3 patch causes loss of nucleolar protein interaction without impairing Mis4 association. IP/mass spectrometry to identify Ssl3-1E2A interactors. Volcano plot compares proteins identified in Ssl3-1E2A-SZZ immunoprecipitates compared to a no tag control. Strains AMfy2094 (*ssl3-1E2A-SZZ*) and AMfy29 (wild type no tag). (D, E) Rad21-6HA ChIP-Seq at centromeres (D) and a chromosomal arm region (E) in wild type (AMfy638) and *dnt1Δ* (AMfy1359) cells.

**Figure S4.**
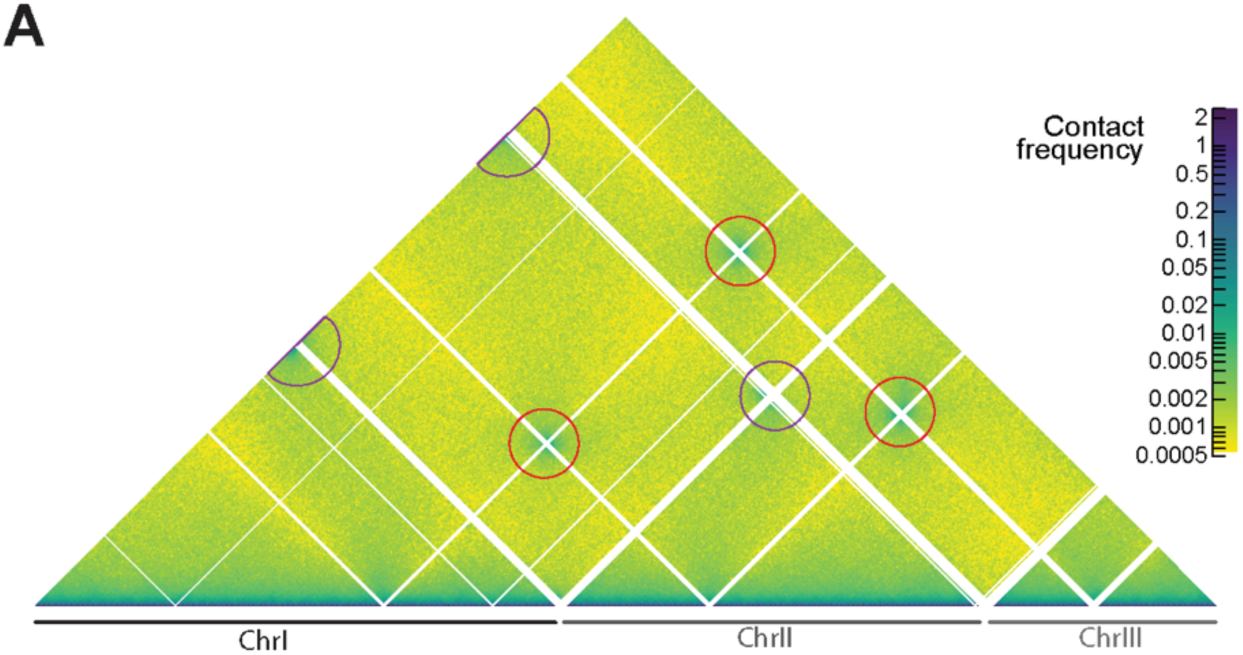
**Chromosomal architecture in *S. pombe.*** (A) Whole genome view of MicroC-XL matrix for wild type (AMfy29) highlighting centromere (red circles) and telomere (purple circles) clustering. Notably, interactions with Chromosome III telomeres are not observed, most likely because the Chromosome III sub-telomeric sequences are missing from the reference genome, resulting from their position between the unsequenced portion of the rDNA repeats and the non-sequenced telomeric sequences (Carme *et al*, 2026).

**Fig S5.**
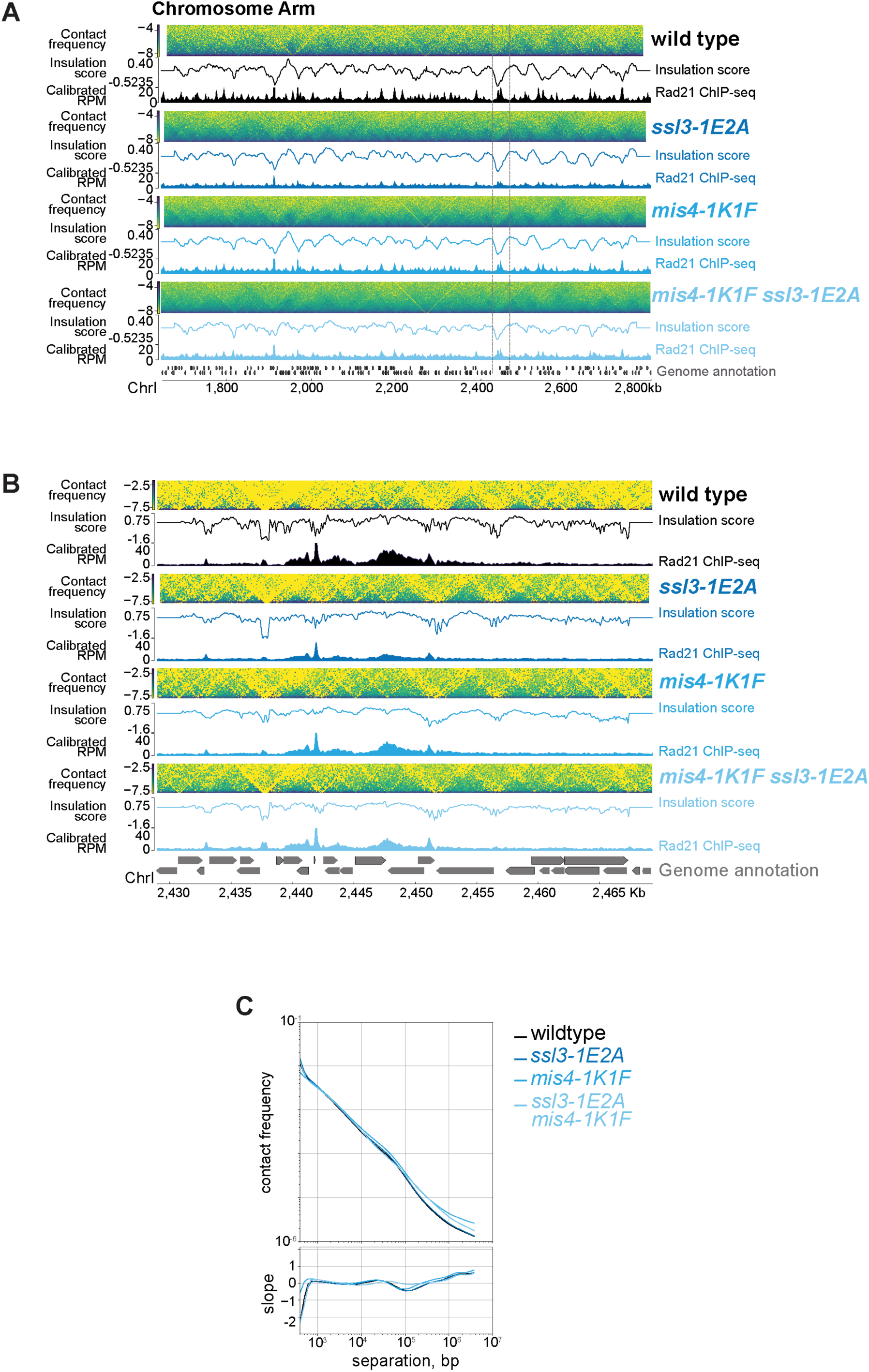
Effect of Mis4-1K1F and Ssl3-1E2A mutations on chromosome domain architecture. (A-B) Bypassing the need for Ssl3 activation of Mis4 broadens chromatin domains without impairing domain boundaries. MicroC-XL matrices at a ∼135kb region of Chromosome I (grey lines denote the region shown in B) (A) and a ∼1.2Mb region of Chromosome I (B), ChIP-seq profiles showing Rad21 enrichment. Contact probability curves and their derivatives for wild type (AMfy29), *ssl3-1E2A* (AMfy1922), *mis4-E907K L1027F* (AMfy2380), *ssl3-1E2A mis4-E907K L1027F* (AMfy2254) made excluding rDNA and pericentromere sequences (C).

**Figure S6.**
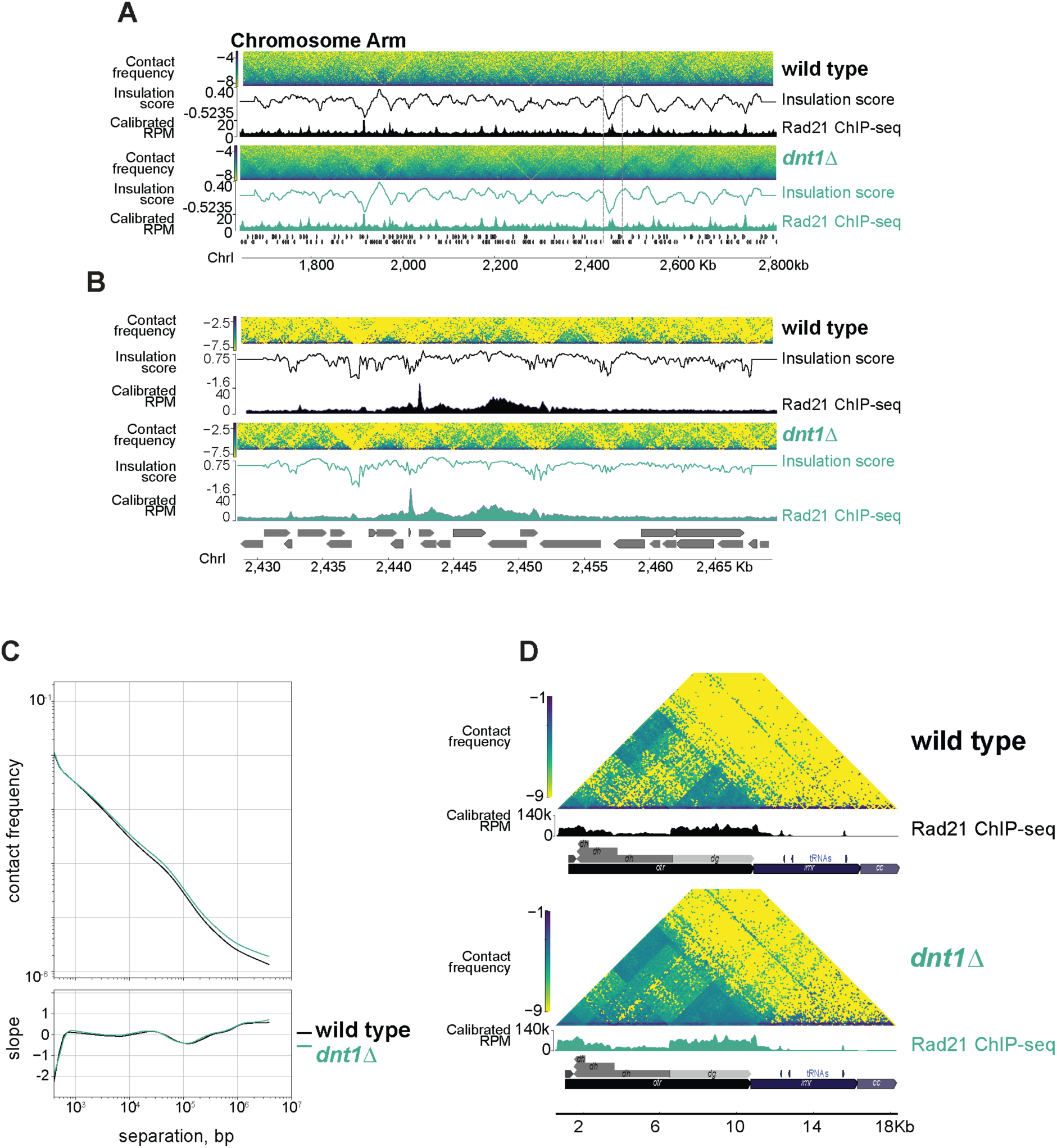
Effect of Dnt1 loss on organisation of arm and pericentromeric chromatin. (A-D) Structure of chromosome arms is unaffected in *dnt1Δ.* (A) Pileups of MicroC-XL matrices from Fig5B at common arm features. (B) Structure of a ∼1.2Mb region of Chromosome I, grey lines denote the zoomed in region shown in (C). (D) contact probability curves and their derivatives. (E-F) Increased levels of cohesin in *dnt1Δ* alters pericentromeres organisation; probability curves and their derivatives indicate increased chromatin contacts (E) which can also be observed on contact matrices (F). Contact matrices are aggregates of data from all three centromeres mapped to the left side of centromere I, genome annotation denotes variable lengths of *dh* repeats.

